# Regulation of Store-Operated Ca^2+^ Entry by IP_3_ Receptors Independent of Their Ability to Release Ca^2+^

**DOI:** 10.1101/2022.04.12.488111

**Authors:** Pragnya Chakraborty, Bipan Kumar Deb, Vikas Arige, Thasneem Musthafa, Sundeep Malik, David I Yule, Colin W. Taylor, Gaiti Hasan

## Abstract

Loss of endoplasmic reticular (ER) Ca^2+^ activates store-operated Ca^2+^ entry (SOCE) by causing the ER localized Ca^2+^ sensor STIM to unfurl domains that activate Orai channels in the plasma membrane at membrane contact sites (MCS). Here we demonstrate a novel mechanism by which the inositol 1,4,5 trisphosphate receptor (IP_3_R), an ER-localized IP_3_-gated Ca^2+^ channel, regulates neuronal SOCE. In human neurons, SOCE evoked by pharmacological depletion of ER-Ca^2+^ is attenuated by loss of IP_3_Rs, and restored by expression of IP_3_Rs even when they cannot release Ca^2+^, but only if the IP_3_Rs can bind IP_3_. Imaging studies demonstrate that IP_3_Rs enhance association of STIM1 with Orai1 in neuronal cells with empty stores; this requires an IP_3_-binding site, but not a pore. Convergent regulation by IP_3_Rs, may tune neuronal SOCE to respond selectively to receptors that generate IP_3_.

## INTRODUCTION

The activities of all eukaryotic cells are regulated by increases in cytosolic free Ca^2+^ concentration ([Ca^2+^]_c_), which are almost invariably evoked by the opening of Ca^2+^- permeable ion channels in biological membranes. The presence of these Ca^2+^ channels within the plasma membrane (PM) and the membranes of intracellular Ca^2+^ stores, most notably the endoplasmic reticulum (ER), allows cells to use both intracellular and extracellular sources of Ca^2+^ to evoke Ca^2+^ signals. In animal cells, the most widely expressed Ca^2+^ signaling sequence links extracellular stimuli, through their specific receptors and activation of phospholipase C, to formation of inositol 1,4,5-trisphosphate (IP_3_), which then stimulates Ca^2+^ release from the ER through IP_3_ receptors (IP_3_R) (Foskett *et al*., 2007; Prole and Taylor, 2019). IP_3_Rs occupy a central role in Ca^2+^ signaling by releasing Ca^2+^ from the ER. IP_3_Rs thereby elicit cytosolic Ca^2+^ signals, and by depleting the ER of Ca^2+^ they initiate a sequence that leads to activation of store-operated Ca^2+^ entry (SOCE) across the PM (Putney, 1986; Thillaiappan *et al*., 2019). SOCE occurs when loss of Ca^2+^ from the ER causes Ca^2+^ to dissociate from the luminal Ca^2+^-binding sites of an integral ER protein, stromal interaction molecule 1 (STIM1). STIM1 then unfolds its cytosolic domains to expose a region that binds directly to a Ca^2+^ channel within the PM, Orai, causing it to open and Ca^2+^ to flow into the cell across the PM (Parekh and Putney, 2005; Prakriya and Lewis, 2015; Lewis, 2020). The interactions between STIM1 and Orai occur across a narrow gap between the ER and PM, a membrane contact site (MCS), where STIM1 puncta trap Orai channels. While STIM1 and Orai are undoubtedly the core components of SOCE, many additional proteins modulate their interactions (Rosado, Jenner and Sage, 2000; Palty *et al*., 2012; Deb, Pathak and Hasan, 2016; Srivats *et al*., 2016) and other proteins contribute by regulating the assembly of MCS (Chang *et al*., 2013; Giordano *et al*., 2013; Kang *et al*., 2019).

It is accepted that IP_3_-evoked Ca^2+^ release from the ER through IP_3_Rs is the usual means by which extracellular stimuli evoke SOCE. Here, the role of the IP_3_R is widely assumed to be restricted to its ability to mediate Ca^2+^ release from the ER and thereby activate STIM1. Evidence from *Drosophila*, where we suggested an additional role for IP_3_Rs in regulating SOCE (Agrawal et al., 2010; Chakraborty et al., 2016), motivated the present study, wherein we examined the contribution of IP_3_Rs to SOCE in mammalian neurons. We show that in addition to their ability to activate STIM1 by evoking ER Ca^2+^ release, IP_3_Rs also facilitate interactions between active STIM1 and Orai1. This additional role for IP_3_Rs, which is regulated by IP_3_ but does not require a functional pore, reveals an unexpected link between IP_3_, IP_3_Rs and Ca^2+^ signaling that is not mediated by IP_3_-evoked Ca^2+^ release. We speculate that dual regulation of SOCE by IP_3_Rs may allow Ca^2+^ release evoked by IP_3_ to be preferentially coupled to SOCE.

## RESULTS

### Loss of IP_3_R1 Attenuates SOCE in Human Neural Stem Cells and Neurons

We investigated the effects of IP_3_Rs on SOCE by measuring [Ca^2+^]_c_ in human neural stem cells and neurons prepared from embryonic stem cells. Human neural progenitor cells (hNPCs) were derived from H9 embryonic stem cells using small molecules that mimic cues provided during human brain development (Gopurappilly *et al*., 2018). We confirmed that hNPCs express canonical markers of neural stem cells (**Figure 1A**) and that IP_3_R1 is the predominant IP_3_R subtype (GEO accession no. GSE109111) (Gopurappilly *et al*., 2018). An inducible lentiviral shRNA-miR construct targeting IP_3_R1 reduced IP_3_R1 expression by 93 ± 0.4% relative to a non-silencing (NS) construct (**Figures 1B and 1C**). Carbachol stimulates muscarinic acetylcholine receptors, which are expressed at low levels in hNPCs (Gopurappilly *et al*., 2018). In Ca^2+^-free medium, carbachol evoked an increase in [Ca^2+^]_c_ in about 10% of hNPCs, consistent with it stimulating Ca^2+^ release from the ER through IP_3_Rs. Restoration of extracellular Ca^2+^ then evoked an increase in [Ca^2+^]_c_ in all cells that responded to carbachol. Both carbachol-evoked Ca^2+^ release and SOCE were abolished in hNPCs expressing IP_3_R1-shRNA, confirming the effectiveness of the IP_3_R1 knockdown (**Figure 1-figure supplement 1A-C**).

**Figure 1.**
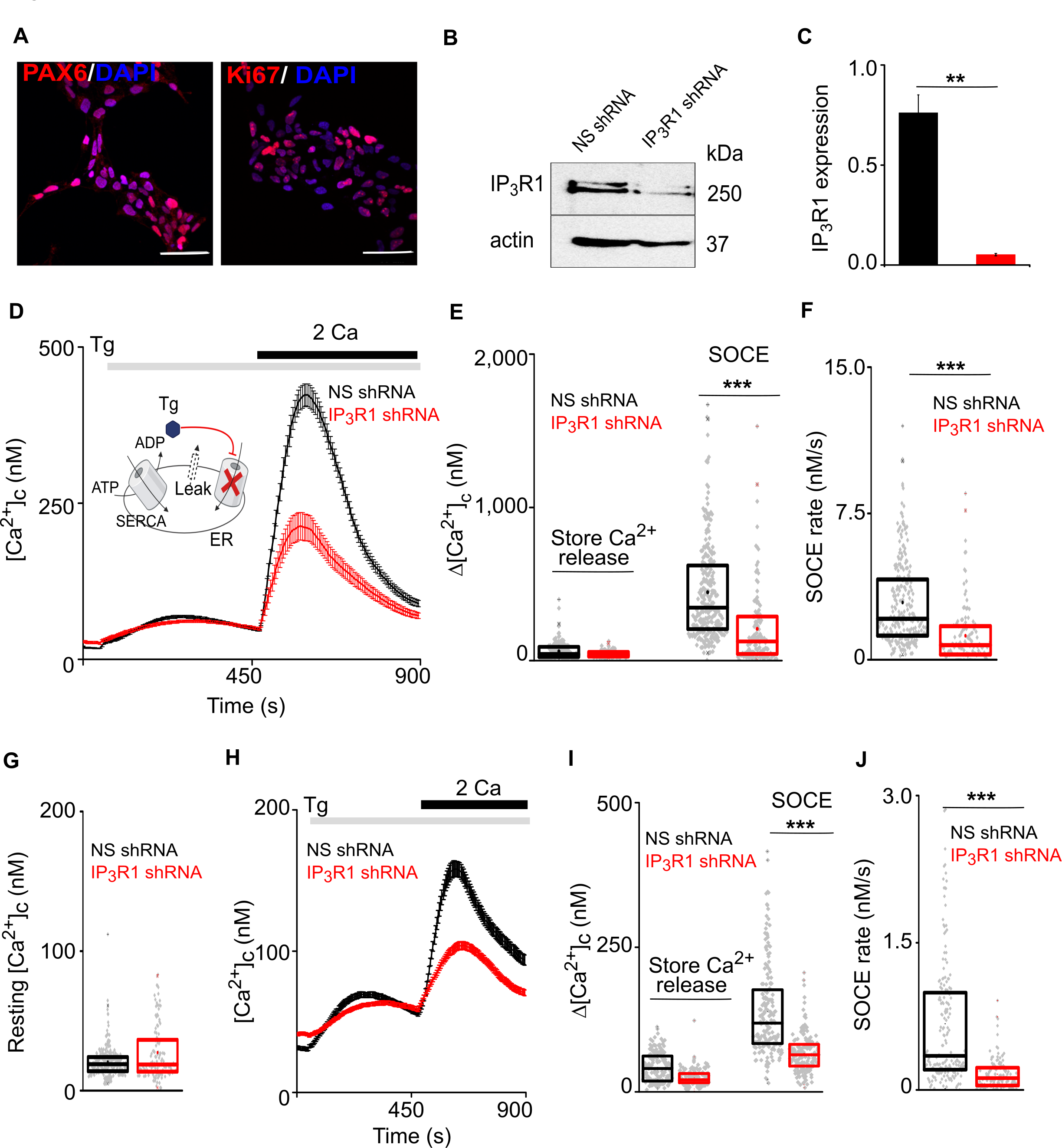
Loss of IP_3_R1 Attenuates SOCE in Human Neural Stem Cells. **(A)**Confocal images of hNPCs (passage 6) stained for DAPI and neural stem cell proteins: Pax6 and Ki67 (proliferation marker). Scale bars, 50 μm. **(B)** WB for IP_3_R1 of hNPCs expressing non-silencing (NS) or IP_3_R1-shRNA. **(C)** Summary results (mean ± s.d., *n* = 3) show IP_3_R1 expression relative to actin. \*\**P* < 0.01, Student’s *t*-test with unequal variances. **(D)** Changes in [Ca^2+^]_c_ evoked by thapsigargin (Tg, 10 µM) in Ca^2+^-free HBSS and then restoration of extracellular Ca^2+^ (2 mM) in hNPCs expressing NS or IP_3_R1- shRNA. Mean ± s.e.m. from 3 independent experiments, each with 4 replicates that together included 100-254 cells. Inset shows the target of Tg. (**E-G**) Summary results (individual cells, median (bar), 25^th^ and 75^th^ percentiles (box) and mean (circle)) show Ca^2+^ signals evoked by Tg or Ca^2+^ restoration (**E**), rate of Ca^2+^ entry (**F**) and resting [Ca^2+^]_c_ (**G**). \*\*\**P* < 0.001, Mann-Whitney U-test. (**H**) Changes in [Ca^2+^]_c_ evoked by Tg (10 µM) in Ca^2+^-free HBSS and after restoring extracellular Ca^2+^ (2 mM) in neurons (differentiated hNPCs) expressing NS or IP_3_R1- shRNA. Mean ± s.e.m. from 3 experiments with ∼200 cells. (**I,J**) Summary results (presented as in E-G) show Ca^2+^ signals evoked by Tg or Ca^2+^ restoration (**I**) and rate of Ca^2+^ entry (**J**). \*\*\**P* < 0.001. Mann-Whitney U-test. See also **Figure 1-figure supplement 1**. **Source data in Figure 1 source data file**

Thapsigargin, a selective and irreversible inhibitor of the ER Ca^2+^ pump (sarcoplasmic/endoplasmic reticulum Ca^2+^-ATPase, SERCA), was used to deplete the ER of Ca^2+^ and thereby activate SOCE (**Figure 1D**) (Parekh and Putney, 2005). Restoration of extracellular Ca^2+^ to thapsigargin-treated hNPCs evoked a large increase in [Ca^2+^]_c_, reflecting the activity of SOCE (**Figure 1D**). The maximal amplitude and rate of SOCE were significantly reduced in cells lacking IP_3_R1, but the resting [Ca^2+^]_c_ and thapsigargin-evoked Ca^2+^ release were unaffected (**Figures 1D-1F** and **Figure 1-figure supplement 1D and 1E**). STIM1 and Orai1 expression were also unaltered in hNPC lacking IP_3_R1 (**Figure 1-figure supplement 1G**). After spontaneous differentiation of hNPC, cells expressed markers typical of mature neurons, and the cells responded to depolarization with an increase in [Ca^2+^]_c_ (**Figure 1-figure supplement 1F and Figure 1-figure supplement 1H-J**). Thapsigargin evoked SOCE in these differentiated neurons; and expression of IP_3_R1-shRNA significantly reduced the SOCE response without affecting depolarization-evoked Ca^2+^ signals (**Figures 1H-1J** and **Figure 1-figure supplement 1H-1L**).

### Loss of IP_3_R1 Attenuates SOCE in Human Neuroblastoma Cells

IP_3_Rs link physiological stimuli that evoke Ca^2+^ release from the ER to SOCE, but the contribution of IP_3_Rs is thought to be limited to their ability to deplete the ER of Ca^2+^. We have reported that in *Drosophila* neurons there is an additional requirement for IP_3_Rs independent of ER Ca^2+^ release (Venkiteswaran and Hasan, 2009; Agrawal *et al*., 2010; Chakraborty *et al*., 2016). Our results with hNPCs and stem cell-derived neurons suggest a similar requirement for IP_3_Rs in regulating SOCE in mammalian neurons. To explore the mechanisms underlying this additional role for IP_3_Rs, we turned to a more tractable cell line, SH-SY5Y cells. These cells are derived from a human neuroblastoma; they exhibit many neuronal characteristics (Agholme *et al*., 2010); they express M3 muscarinic acetylcholine receptors that evoke IP_3_-mediated Ca^2+^ release and SOCE (Grudt, Usowicz and Henderson, 1996); and they express predominantly IP_3_R1 (Wojcikiewicz, 1995; Tovey *et al*., 2001), with detectable IP_3_R3, but no IP_3_R2 (**Figure 2A**). We used inducible expression of IP_3_R1-shRNA to significantly reduce IP_3_R1 expression (by 74 ± 1.2%), without affecting IP_3_R3 (**Figures 2A and 2B**). As expected, carbachol-evoked Ca^2+^ signals in individual SH-SY5Y cells were heterogenous and the carbachol-evoked Ca^2+^ release was significantly reduced by knockdown of IP_3_R1 (**Figures 2C and 2D** and **Figure 2 - figure supplement 1A and 1B**). Thapsigargin evoked SOCE in SH-SY5Y cells (Grudt, Usowicz and Henderson, 1996), and it was significantly attenuated after knockdown of IP_3_R1 without affecting resting [Ca^2+^]_c_, the Ca^2+^ release evoked by thapsigargin or expression of STIM1 and Orai1 (**Figures 2E-2G** and **Figure 2 - figure supplement 1C-1E**).

**Figure 2.**
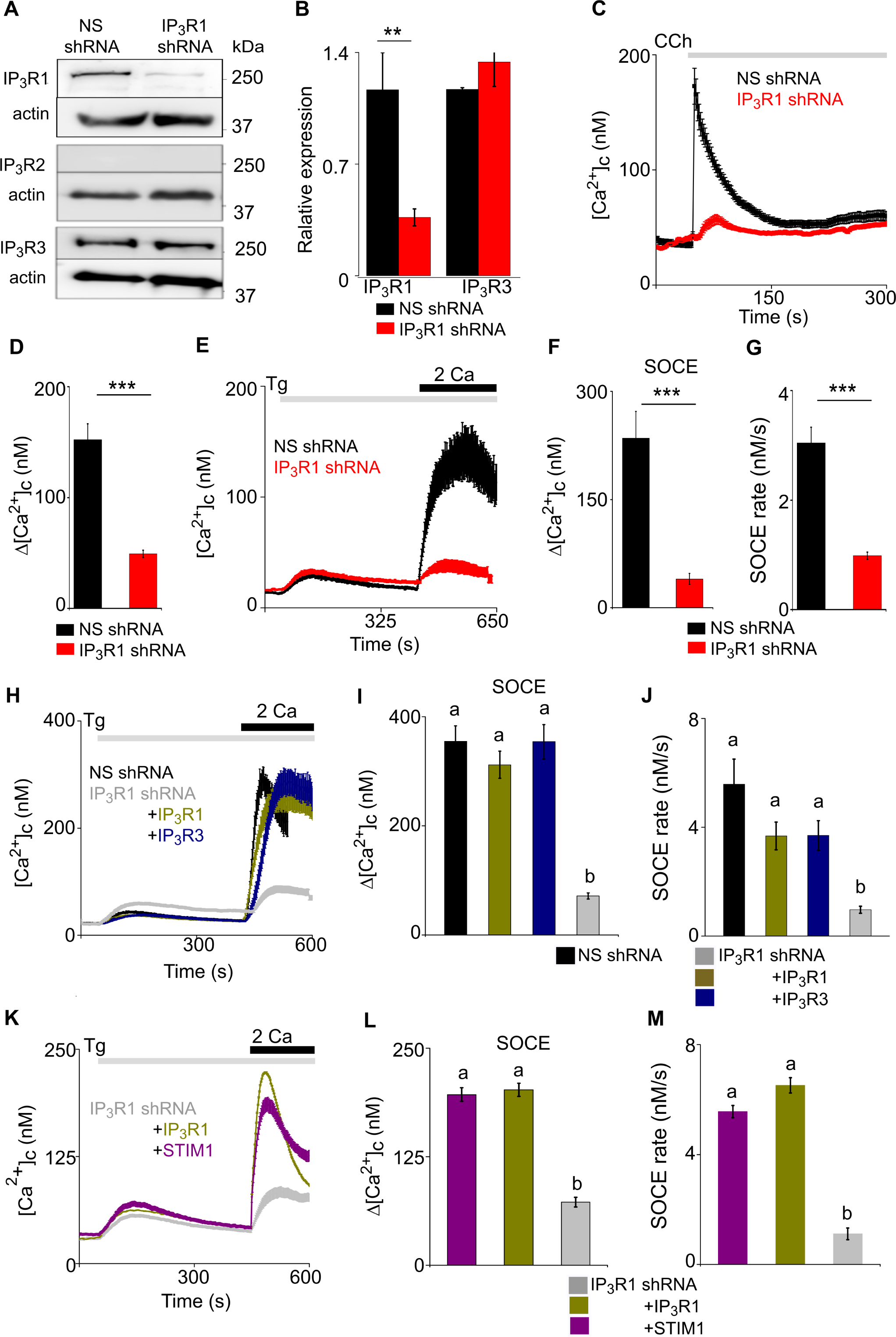
Loss of IP_3_R1 Attenuates SOCE in SH-SY5Y Cells. **(A)**WB for IP_3_R1-3 of SH-SY5Y cells expressing non-silencing (NS) or IP_3_R1-shRNA. **(B)** Summary results (mean ± s.d., *n* = 4) show IP_3_R expression relative to actin normalized to control NS cells. \*\**P* < 0.01, Student’s *t*-test with unequal variances. **(C)** Ca^2+^ signals evoked by carbachol (CCh, 3 µM) in SH-SY5Y cells expressing NS or IP_3_R1-shRNA. Mean ± s.e.m. from 3 experiments with 70-90 cells. **(D)** Summary results show peak changes in [Ca^2+^]_c_ (Δ[Ca^2+^]_c_) evoked by CCh. \*\*\**P* < 0.001, Mann-Whitney U-test. **(E)** Ca^2+^ signals evoked by thapsigargin (Tg, 10 µM) in Ca^2+^-free HBSS and then after restoration of extracellular Ca^2+^ (2 mM) in cells expressing NS or IP_3_R1-shRNA. Mean ± s.e.m. from 3 experiments with ∼50 cells. (**F, G**) Summary results (individual cells, mean ± s.e.m., *n* = 3, ∼50 cells) show peak changes in [Ca^2+^]_c_ evoked by Ca^2+^ restoration (Δ[Ca^2+^]_c_) (**F**) and rate of Ca^2+^ entry (**G**). \*\*\**P* < 0.001, Mann-Whitney U-test. (**H**) Ca^2+^ signals evoked by Tg and then Ca^2+^ restoration in cells expressing NS-shRNA, or IP_3_R1-shRNA alone or with IP_3_R1 or IP_3_R3. Traces show mean ± s.e.m. (50-115 cells from 3 experiments). (**I, J**) Summary results (mean ± s.e.m, 50-115 cells from 3 experiments) show peak increases in [Ca^2+^]_c_ (Δ[Ca^2+^]_c_) evoked by Ca^2+^ restoration (**I**) and rates of Ca^2+^ entry (**J**) evoked by restoring extracellular Ca^2+^. (**K**) Effects of thapsigargin (Tg, 10 µM) in Ca^2+^-free HBSS and then after Ca^2+^ restoration (2 mM) in cells expressing IP_3_R1-shRNA alone or with IP_3_R1 or mCh-STIM1. Traces show mean ± s.e.m. (100-150 cells from 3 experiments). (**L, M**) Summary results (mean ± s.e.m.) show peak increase in [Ca^2+^]_c_ after Ca^2+^ restoration (Δ[Ca^2+^]_c_) (**L**) and rate of Ca^2+^ entry (**M**). Different letters indicate significant differences (panels I, J, L, M), *P* < 0.001, one-way ANOVA with pair-wise Tukey’s test. See alsog **Figure 2 with 3 supplements (Figure 2 - figure supplements 1 - 3)**. **Source data in Figure 2 source data file**

We also used CRISPR/Cas9n and Cas9 to disrupt one or both copies of the IP_3_R1 gene, subsequently referred to as IKO (one copy knockout) and IKO null (both copies knocked out) in SH-SY5Y cells. IP_3_R1 expression was absent in the IKO null (**Figure 2 - figure supplement 1F**) whereas expression of STIM1, STIM2 and Orai1 were unperturbed (**Figure 2 - figure supplement 1G**). Carbachol-evoked Ca^2+^ release and thapisgargin-evoked SOCE were significantly reduced (**Figure 2 - figure supplement 1H-1J**). Since the IKO null cells were fragile and grew slowly, we examined SOCE in SH-SY5Y cells with disruption of one copy of the IP_3_R1 gene. In the IKO cells, IP_3_R1 expression, carbachol-evoked Ca^2+^ signals and thapsigargin- evoked SOCE were all reduced (**Figure 2- figure supplement 1K-1Q**).

These observations, which replicate those from hNPCs and neurons (**Figure 1**), vindicate our use of SH-SY5Y cells to explore the mechanisms linking IP_3_Rs to SOCE in human neurons.

Expression of IP_3_R1 or IP_3_R3 in SH-SY5Y cells expressing IP_3_R1-shRNA restored both carbachol-evoked Ca^2+^ release and thapsigargin-evoked SOCE without affecting resting [Ca^2+^]_c_ or thapsigargin-evoked Ca^2+^ release (**Figures 2H-2J** and **Figure 2 - figure supplement 2A-2D**). Over-expression of STIM1 in cells expressing NS-shRNA had no effect on SOCE (**Figure 2 - figure supplement 2E and 2F**), but it restored thapsigargin-evoked SOCE in cells expressing IP_3_R1-shRNA, without affecting resting [Ca^2+^]_c_ or thapsigargin-evoked Ca^2+^ release (**Figures 2K-2M**). We conclude that IP_3_Rs are required for optimal SOCE, but they are not essential because additional STIM1 can replace the need for IP_3_Rs (**Figure 3A**).

**Figure 3.**
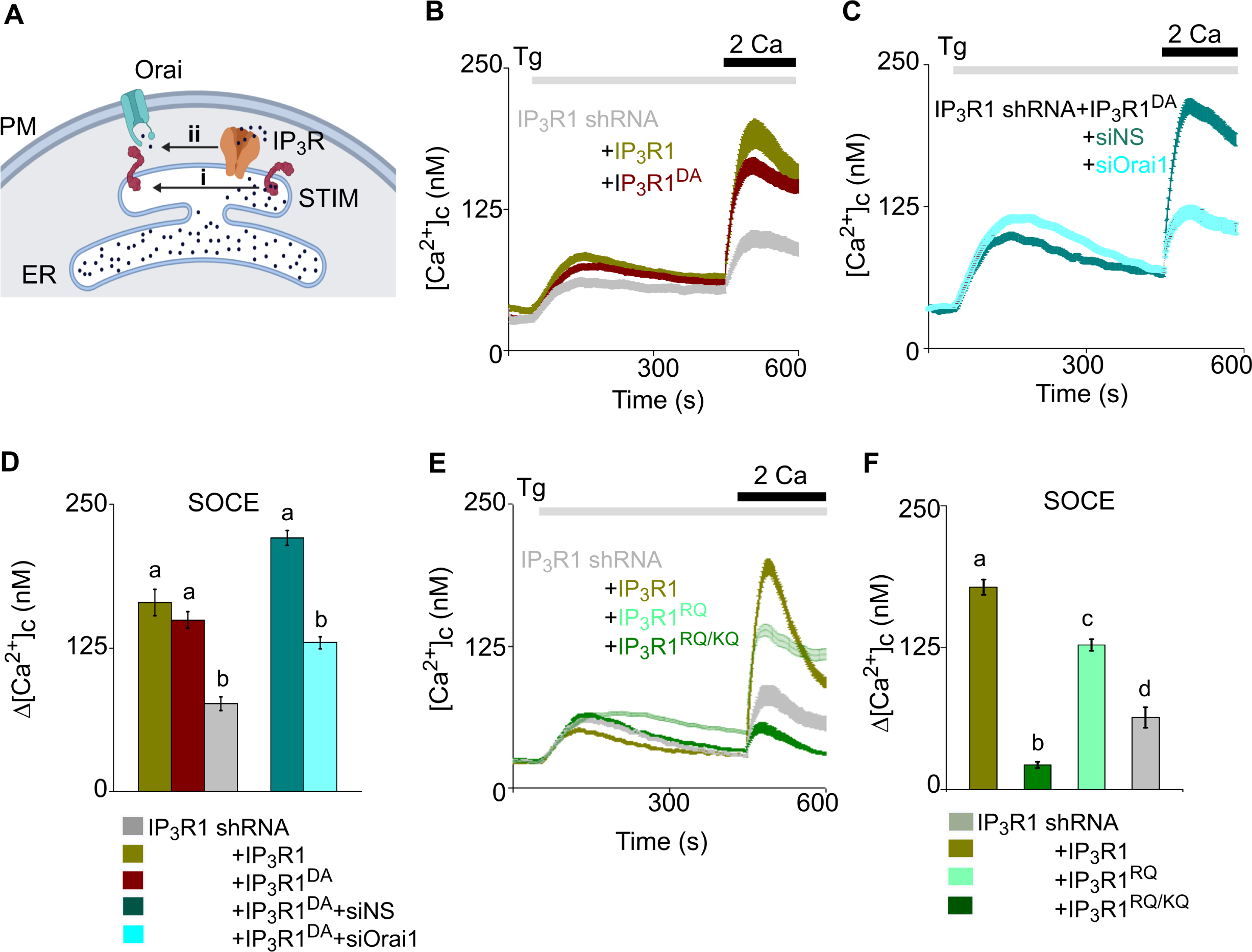
Regulation of SOCE by IP_3_R Requires IP_3_ Binding but Not a Functional Pore in SH-SY5Y cells. **(A)**SOCE is activated when loss of Ca^2+^ from the ER through IP_3_Rs activates STIM1 (i). Our results suggest an additional role for IP_3_Rs (ii). **(B)** SH-SY5Y cells expressing IP_3_R1-shRNA alone or with IP_3_R1 or IP_3_R1^DA^ were stimulated with thapsigargin (Tg, 1 µM) in Ca^2+^-free HBSS before restoring extracellular Ca^2+^ (2 mM). Traces show mean ± s.e.m, for 100-150 cells from 3 experiments. **(C)** Cells expressing IP_3_R1-shRNA and IP_3_R1^DA^ were treated with NS-siRNA or Orai1- siRNA before measuring Tg-evoked Ca^2+^ entry. Traces show mean ± s.e.m. for 85- 100 cells from 3 experiments. **(D)** Summary results (mean ± s.e.m.) show peak increases in [Ca^2+^]_c_ (Δ[Ca^2+^]_c_) evoked by Ca^2+^ restoration. **(E)** Tg-evoked Ca^2+^ entry in cells expressing IP_3_R1-shRNA with IP_3_R1, IP_3_R1^RQ^ or IP_3_R1^RQ/KQ^. Traces show mean ± s.e.m, for 90-150 cells from 3 experiments. **(F)** Summary results (mean ± s.e.m.) show peak increases in [Ca^2+^]_c_ (Δ[Ca^2+^]_c_) evoked by Ca^2+^ restoration. Different letter codes (panels D, F) indicate significantly different values, *P* < 0.001, for multiple comparison one-way ANOVA and pair-wise Tukey’s test and for two genotype comparison Mann Whitney U-test. See also **Figure 3 - figure supplement 1**. **Source data in Figure 3 source data file**

It has been reported that SOCE is unaffected by loss of IP_3_R in non-neuronal cells (Ma *et al*., 2001; Chakraborty *et al*., 2016). Consistent with these observations, the SOCE evoked in HEK cells by stores emptied fully by treatment with thapsigargin was unaffected by expression of IP_3_R1 shRNA (**Figure 2 - figure supplement 3A-3C**) or by knockout of all three IP_3_R subtypes using CRISPR/cas9 (HEK-TKO cells) (**Figure 2 - figure supplement 3D and 3E**). The association of STIM1 with Orai1 in wild type HEK cells and HEK TKO cells after thapsigargin-evoked store depletion also appeared identical as tested by a proximity ligation assay (PLA, described further in **Figure 5) (Figure 2 - figure supplement 3F**). Neuronal and non-neuronal cells may, therefore, differ in the contribution of IP_3_R to SOCE. We return to this point later.

### Binding of IP_3_ to IP_3_R Without a Functional Pore Stimulates SOCE

IP_3_Rs are large tetrameric channels that open when they bind IP_3_ and Ca^2+^, but they also associate with many other proteins (Prole and Taylor, 2019), and many IP_3_Rs within cells appear not to release Ca^2+^ (Thillaiappan *et al*., 2019). A point mutation (D2550A, IP_3_R1^D/A^) within the IP_3_R1 pore prevents it from conducting Ca^2+^ (Dellis *et al*., 2008). As expected, expression of IP_3_R1^D/A^ in cells lacking IP_3_R1 failed to rescue carbachol-evoked Ca^2+^ release, but it unexpectedly restored thapsigargin-evoked SOCE (**Figures 3B-3D** and **Figure 3 - figure supplement 1A-E**). We confirmed that rescue of thapsigargin-evoked Ca^2+^ entry by this pore-dead IP_3_R was mediated by a conventional SOCE pathway by demonstrating that it was substantially attenuated by siRNA-mediated knockdown of Orai1 (**Figures 3C and 3D** and **Figure 3 - figure supplement 1F-1H**).

Activation of IP_3_Rs is initiated by IP_3_ binding to the N-terminal IP_3_-binding core of each IP_3_R subunit (Prole and Taylor, 2019). Mutation of two conserved phosphate- coordinating residues in the α-domain of the binding core (R568Q and K569Q of IP_3_R1, IP_3_R1^RQ/KQ^) almost abolishes IP_3_ binding (Yoshikawa *et al*., 1996; Iwai *et al*., 2007), while mutation of a single residue (R568Q, IP_3_R1^RQ^) reduces the IP_3_ affinity by ∼10-fold (Dellis *et al*., 2008). Expression of rat IP_3_R1^RQ/KQ^ rescued neither carbachol- evoked Ca^2+^ release nor thapsigargin-evoked SOCE in cells lacking IP_3_R1 (**Figures 3E and 3F** and **Figure 3 - figure supplement 1C and 1I**). However, expression of IP_3_R1^RQ^ substantially rescued thapsigargin-evoked SOCE (**Figures 3E and 3F** and **Figure 3 - figure supplement 1J**). Expression of an N-terminal fragment of rat IP_3_R (IP_3_R1^1-604^), to which IP_3_ binds normally (Iwai *et al*., 2007), failed to rescue thapsigargin-evoked SOCE (**Figure 3 - figure supplement 1K and 1L**). These results establish that a functional IP_3_-binding site within a full-length IP_3_R is required for IP_3_Rs to facilitate thapsigargin-evoked SOCE. Hence in cells with empty Ca^2+^ stores, IP_3_ binding, but not pore-opening, is required for regulation of SOCE by IP_3_Rs. In cells stimulated only with thapsigargin and expressing IP_3_Rs with deficient IP_3_ binding, basal levels of IP_3_ are probably insufficient to meet this need.

We further examined the need for IP_3_ by partially depleting the ER of Ca^2+^ using cyclopiazonic acid (CPA), a reversible inhibitor of SERCA, to allow submaximal activation of SOCE (**Figure 3 - figure supplement 1M and 1N**). Under these conditions, addition of carbachol in Ca^2+^-free HBSS to SH-SY5Y cells expressing IP_3_R1-shRNA caused a small increase in [Ca^2+^]_c_ (**Figures 4A-4C**). In the same cells expressing IP_3_R1^DA^, the carbachol-evoked Ca^2+^ release was indistinguishable from that observed in cells without IP_3_R^DA^ (**Figures 4B and 4C**), indicating that the small response was entirely mediated by residual native IP_3_R1 and/or IP_3_R3. Hence, the experiment allows carbachol to stimulate IP_3_ production in cells expressing IP_3_R1^DA^ without causing additional Ca^2+^ release. The key result is that in cells expressing IP_3_R1^DA^, carbachol substantially increased SOCE (**Figures 4A-4C**). Moreover, addition of carbachol to control shRNA expressing SH-SY5Y cells with maximal store depletion (thapsigargin, Tg, 2µM) also resulted in increased SOCE (**Figure 4 – figure supplement 1A**). We conclude that in neuronal cells IP_3_, through IP_3_Rs, regulates coupling of empty stores to SOCE. This is the first example of an IP_3_R mediating a response to IP_3_ that does not require the pore of the channel.

**Figure 4.**
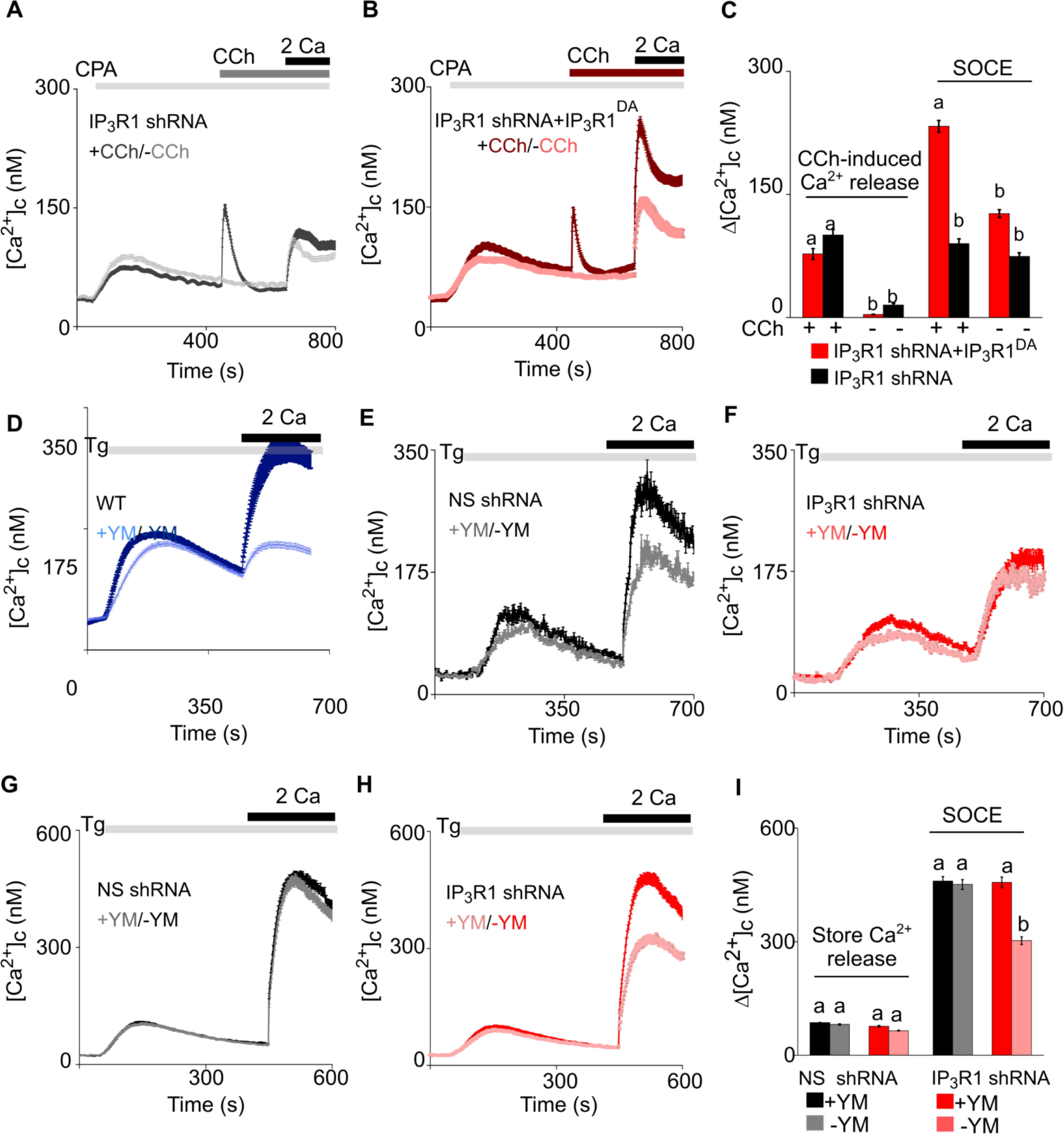
Receptor-Regulated IP_3_ Production Stimulates SOCE in Cells With Empty Ca^2+^ Stores and Expressing Pore-Dead IP_3_R. (**A, B**) SH-SY5Y cells expressing IP_3_R1-shRNA alone (**A**) or with IP_3_R1^DA^ (**B**) were treated with a low concentration of CPA (2 µM) in Ca^2+^-free HBSS to partially deplete the ER of Ca^2+^ and sub-maximally activate SOCE (see **Figures S5M and S5N**). Carbachol (CCh, 1 µM) was then added to stimulate IP_3_ formation through muscarinic receptors, and extracellular Ca^2+^ (2 mM) was then restored. Traces (mean ± s.e.m of 68-130 cells from 3 experiments) show responses with and without the CCh addition. (**C**) Summary results show the peak increases in [Ca^2+^]_c_ (Δ[Ca^2+^]_c_) after addition of CCh (CCh-induced Ca^2+^ release) and then after restoring extracellular Ca^2+^ (SOCE). (**D-F**) SH-SY5Y cells wild type (WT) (**D**) and expressing NS-shRNA (**E**) or IP_3_R1- shRNA (**F**) were treated with YM-254890 (YM, 1 µM, 5 min) in Ca^2+^-free HBSS to inhibit Gαq and then with thapsigargin (Tg, 1 µM) before restoring extracellular Ca^2+^ (2 mM). Traces show mean ± s.e.m of ∼120 cells from 3 experiments. . (**G-I**) Similar analyses of HEK cells. Summary results (mean ± s.e.m, 50-100 cells from 3 experiments) are shown in (I). Different letter codes (panels C and I) indicate significantly different values within the store Ca^2+^ release or SOCE groups, *P* < 0.001, one-way ANOVA and pair-wise Tukey’s test. See also **Figure 4 - figure supplement 1**. **Source data in Figure 4 source data file**

G-protein-coupled receptors are linked to IP_3_ formation through the G-protein Gq, which stimulates phospholipase C β (PLC β). We used YM-254890 to inhibit Gq (Kostenis, Pfeil and Annala, 2020; Patt *et al*., 2021). As expected, addition of YM- 254890 to wild type (WT) or NS-shRNA transfected SH-SY5Y cells abolished the Ca^2+^ signals evoked by carbachol **(Figure 4 - figure supplement 1C)**, but it also reduced the maximal amplitude and rate of thapsigargin-evoked SOCE (**Figures 4D-4E** and **Figure 3 - figure supplement 1O**). YM-254890 had no effect on the residual thapsigargin-evoked SOCE in SH-SY5Y cells expressing IP_3_R1-shRNA (**Figure 4F** and **Figure 3 - figure supplement 1O**). The latter result is important because it demonstrates that the inhibition of SOCE in cells with functional IP_3_Rs is not an off- target effect causing a direct inhibition of SOCE.

In wild type or HEK-TKO (lacking all three IP_3_Rs) cells, YM-254890 had no effect on thapsigargin-evoked SOCE, but it did inhibit SOCE in HEK cells lacking only IP_3_R1 (**Figures 4G-4I** and **Figure 4 - figure supplement 1D-1G**). These results suggest that in HEK cells, which normally express all three IP_3_R subtypes (Mataragka & Taylor, 2018), neither loss of IP_3_R1 nor inhibition of Gαq is sufficient on its own to inhibit thapsigargin-evoked SOCE, but when combined there is a synergistic loss of SOCE.

### IP_3_Rs Promote Interaction of STIM1 With Orai1 Within MCS

Our evidence that IP_3_Rs intercept coupling between empty stores and SOCE (**Figure 3A**) prompted us to investigate the coupling of STIM1 with Orai1 across the narrow junctions between ER and PM (Carrasco and Meyer, 2011). An *in situ* proximity ligation assay (PLA) is well suited to analyzing this interaction because it provides a signal when two immunolabelled proteins are within ∼40 nm of each other (Derangère *et al*., 2016), a distance comparable to the dimensions of the junctions wherein STIM1 and Orai1 interact (Poteser *et al*., 2016). We confirmed the specificity of the PLA and demonstrated that it reports increased association of STIM1 with Orai1 after treating SH-SY5Y cells with thapsigargin by measuring the surface area of PLA spots (**Figure 5A** and **Figure 5 - figure supplement 1A-1F**) and not the number, because the latter did not change upon store-depletion (**Figure 5 – figure supplement 1O**). In cells expressing IP_3_R1-shRNA, thapsigargin had no effect on the STIM1-Orai1 interaction reported by PLA, but the interaction was rescued by expression of IP_3_R1 or IP_3_R1^DA^. There was no rescue with IP_3_R1^RQ/KQ^ (**Figures 5B-5E**). WT SH-SY5Y cells that were depleted of basal IP_3_ by treatment with the Gq inhibitor YM-254890, showed significantly reduced STIM1-Orai1 interaction after thapsigargin-evoked depletion of Ca^2+^ stores (**Figure 4 - figure supplement 1B**). The results with PLA exactly mirror those from functional analyses (**Figures 1** to **4**), suggesting that IP_3_ binding to IP_3_R enhances SOCE by facilitating interaction of STIM1 with Orai1 (**Figure 3A**).

**Figure 5.**
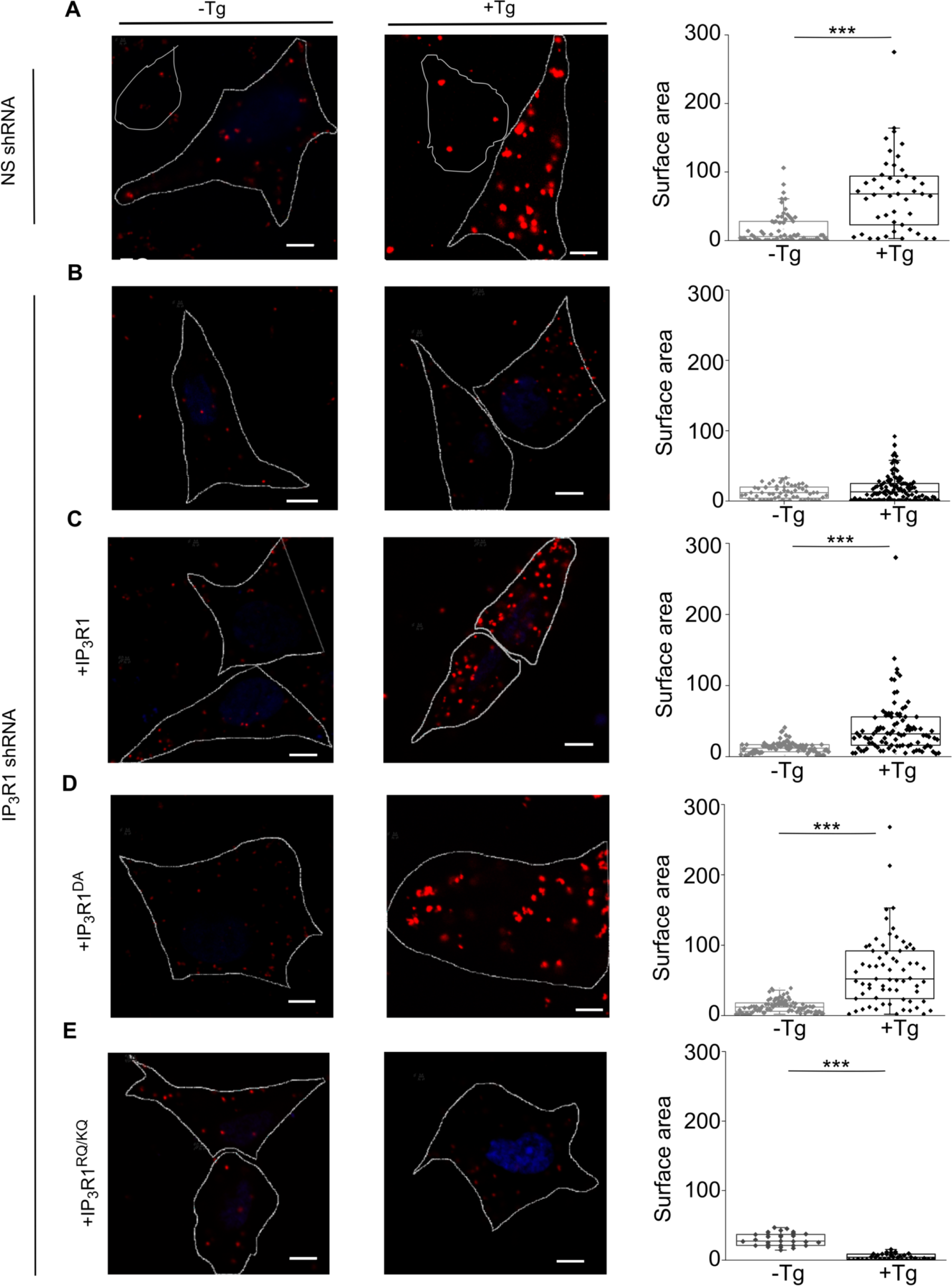
IP_3_Rs Promote Interaction of STIM1 With Orai1. (**A-E**) PLA analyses of interactions between STIM1 and Orai1 in SH-SY5Y cells expressing NS-shRNA (**A**) or IP_3_R1-shRNA alone (**B**) or with IP_3_R1 (**C**), IP_3_R1^DA^ (**D**) or IP_3_R1^RQ/KQ^ (**E**). Confocal images are shown for control cells or after treatment with thapsigargin (Tg, 1 µM) in Ca^2+^-free HBSS. PLA reaction product is red, and nuclei are stained with DAPI (blue). Scale bars, 5 µm. Summary results show the surface area of the PLA spots for 8-10 cells from 2 independent analyses. Individual values, median (bar) and 25^th^ and 75^th^ percentiles (box). \*\*\**P* < 0.001, Student’s *t*-test with unequal variances. See also **Figure 5 - figure supplement 1**. **Source data in Figure 5 source data file**

In independent experiments we tested the effect of fluorescent-tagged and ectopically expressed ligand bound (wild type rat IP_3_R1) and mutant (rat IP_3_R1^RQ/KQ^; **Figure 6- figure supplement 1A)** IP_3_R1 on SOCE dependent STIM1 oligomerization and translocation to ER-PM junctions in SH-SY5Y cells (**Figure 6**). Consistent with previous observations, SOCE did not bring about a measurable change in ER-PM translocation of over-expressed wild type mCherry-IP_3_R1 and mCherry-IP_3_R1^RQ/KQ^ constructs (**Figure 6A-6C**). In agreement with PLA data (**Figure 5**) ER-PM translocation of mVenus-STIM1 upon SOCE induction was reduced significantly in mCherry-IP_3_R1^RQ/KQ^ expressing cells compared to mCherry-IP_3_R1 expressing SH- SY5Y cells (**Figures 6A, 6B, 6D, and 6E**).

**Figure 6.**
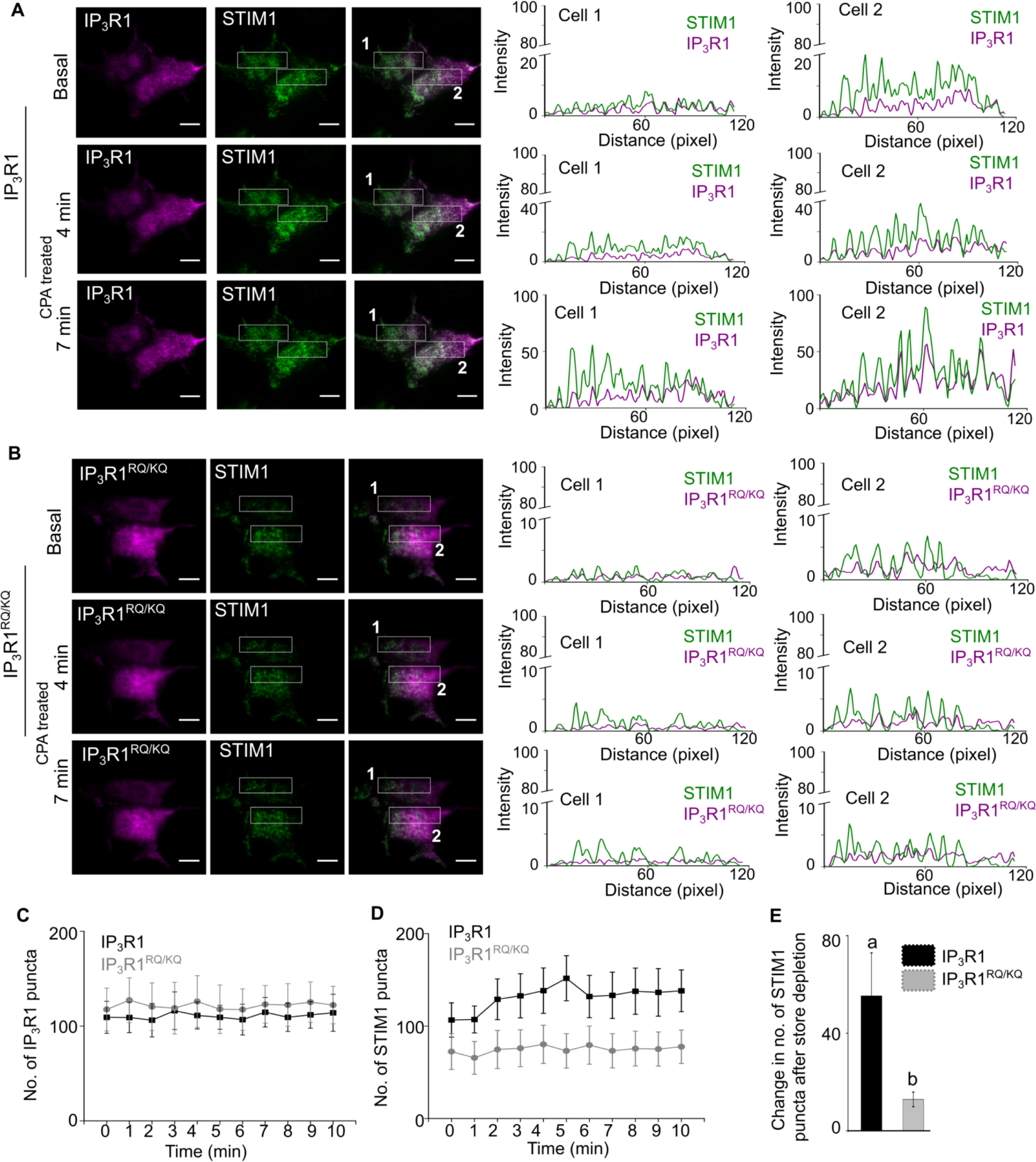
Ligand-bound IP_3_R1 supports SOCE-dependent STIM1 movement to ER-PM contact sites. (**A-B)** Representative TIRF images of mVenus STIM1 co-transfected with either wild type mcherry-rat IP_3_R1 (**A**) or IP_3_R1^RQ/KQ^ (ligand binding mutant), (**B**) in wild type SH- SY5Y cells before (Basal) and after CPA induced store depletion (CPA treated) at 4 min and 7min. On the right are shown RGB profile plots of STIM1 (green) and IP_3_R1, wild type or mutant (magenta) corresponding to the rectangular selections (Cell 1 and Cell 2). Scale bar is 10µm. **(C-D)** Changes in number of IP_3_R1 (**C**) and STIM1 (**D**) puncta upon CPA-induced store depletion over a period of 10 min in the indicated genotypes. Mean ± s.e.m from 7 cells from n=6 independent experiments. **(E)** Summary result (mean ± s.e.m) showing the change in the number of maximum STIM1 puncta formed after CPA-induced store depletion in the indicated genotypes. Mean ± s.e.m. of 7 cells from n=6 independent experiments. Different letters indicate significant differences, P < 0.05, Mann-Whitney U-test. See also **Figure 6 - figure supplement 1**. **Source data in Figure 6 source data file.**

Extended synaptotagmins (E-Syts) are ER proteins that stabilize ER-PM junctions including STIM1-Orai1 MCS (Maléth *et al*., 2014; Kang *et al*., 2019; Woo *et al*., 2020). Over-expression of E-Syt1 in SH-SY5Y cells expressing IP_3_R1-shRNA rescued thapsigargin-evoked Ca^2+^ entry without affecting resting [Ca^2+^]_c_ or thapsigargin-evoked Ca^2+^ release (**Figures 7A-7C**). The rescued Ca^2+^ entry is likely to be mediated by conventional SOCE because it was substantially attenuated by knockdown of STIM1 (**Figures 7D-7F**). Over-expression of E-Syt1 had no effect on SOCE in cells with unperturbed IP_3_Rs (**Figures 7G-7I**). These results suggest that attenuated SOCE after loss of IP_3_Rs can be restored by exaggerating ER-PM MCS.

**Figure 7.**
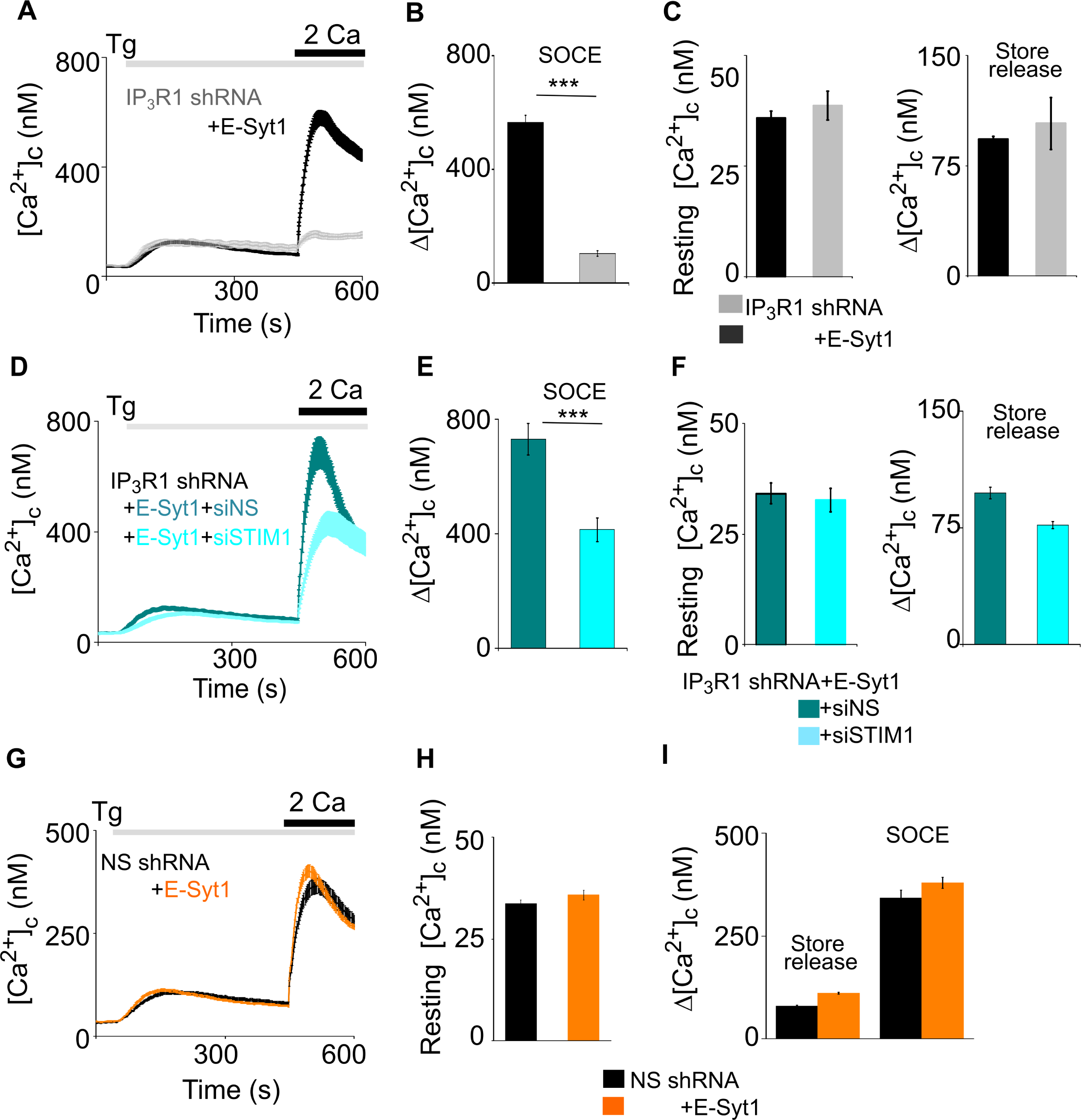
Extended Synaptotagmins Rescue SOCE in Cells Lacking IP_3_R1. **(A)**SH-SY5Y cells expressing IP_3_R1-shRNA alone or with E-Syt1 were stimulated with Tg (1 µM) in Ca^2+^-free HBSS before restoring extracellular Ca^2+^ (2 mM). Traces show mean ± s.e.m, for 20-80 cells from 3 experiments. **(B)** Summary results show Δ[Ca^2+^]_c_ evoked by restoring Ca^2+^ (SOCE). Mean ± s.e.m, \*\*\**P* < 0.001, Mann-Whitney U- test. **(C)** Summary results (mean ± s.e.m, n = 20-80 cells) show resting [Ca^2+^]_c_ (left) and the peak Ca^2+^ signals (Δ[Ca^2+^]_c_) evoked by thapsigargin (Tg, 1 µM) in Ca^2+^-free HBSS for SH-SY5Y cells expressing IP_3_R1-shRNA alone or with human E-Syt1. **(D)** Cells over-expressing E-Syt1 and treated with IP_3_R1-shRNA in combination with either NS or STIM1 siRNA were stimulated with Tg (1 µM) in Ca^2+^-free HBSS before restoration of extracellular Ca^2+^ (2 mM). Mean ± s.e.m. from 3 experiments with 30-40 cells. (**E, F**) Summary results (mean ± s.e.m, n = 30-40 cells) show SOCE evoked by Tg (**E**), resting [Ca^2+^]_c_ and the Tg-evoked Ca^2+^ release from intracellular stores (F). \*\*\**P* < 0.001, Mann-Whitney U- test. **(G)** Similar analyses of cells expressing NS shRNA alone or with human E-Syt1 and then treated with Tg (1 µM) in Ca^2+^-free HBSS before restoring extracellular Ca^2+^ (2 mM). Mean ± s.e.m. from 3 experiments with 115-135 cells. (**H, I**) Summary results (mean ± s.e.m, n = 115-135 cells) show resting [Ca^2+^]_c_ (**H**) and Δ[Ca^2+^]_c_ evoked by Tg (store release) or Ca^2+^ restoration (SOCE) (**I**). No significant difference, Mann Whitney U-test. **Source data in Figure 7 source data file**

## DISCUSSION

After identification of STIM1 and Orai1 as core components of SOCE (Prakriya and Lewis, 2015; Thillaiappan *et al*., 2019), the sole role of IP_3_Rs within the SOCE pathway was assumed to be the release of ER Ca^2+^ that triggers STIM1 activation. The assumption is consistent with evidence that thapsigargin-evoked SOCE can occur in avian (Sugawara *et al*., 1997; Ma *et al*., 2002; Chakraborty *et al*., 2016) and mammalian cells without IP_3_Rs (Prakriya and Lewis, 2001). Although SOCE in mammalian HEK cells was unaffected by loss of IP_3_Rs in our study (**Figure 2-figure supplement 3)**, it was modestly reduced in other studies of mammalian cells (Bartok *et al*., 2019; Yue *et al*., 2020). However, additional complexity is suggested by evidence that SOCE may be reduced in cells without IP_3_Rs (Chakraborty *et al*., 2016; Bartok *et al*., 2019; Yue *et al*., 2020), by observations implicating phospholipase C in SOCE regulation (Rosado, Jenner and Sage, 2000; Broad *et al*., 2001), by evidence that SOCE responds differently to IP_3_Rs activated by different synthetic ligands (Parekh, Riley and Potter, 2002) and by some, albeit conflicting reports (Woodard *et al*., 2010; Santoso, Cebotaru and Guggino, 2011; Béliveau, Lessard and Guillemette, 2014; Sampieri *et al*., 2018; Ahmad *et al*., 2022), that IP_3_Rs may interact with STIM and/or Orai (Woodard *et al*., 2010; Santoso, Cebotaru and Guggino, 2011; Béliveau, Lessard and Guillemette, 2014; Sampieri *et al*., 2018).

We identified two roles for IP_3_Rs in controlling endogenous SOCE in human neurons. As widely reported, IP_3_Rs activate STIM1 by releasing Ca^2+^ from the ER, but they also, and independent of their ability to release Ca^2+^, enhance interactions between active STIM1 and Orai1 (**Figure 8**). The second role for IP_3_Rs can be supplanted by over-expressing other components of the SOCE complex, notably STIM1 or ESyt1 (**Figures 2K-2M** and **Figures 7A and 7B**). It is intriguing that STIM1 (Carrasco and Meyer, 2011; Lewis, 2020), ESyt1 (Giordano *et al*., 2013) and perhaps IP_3_Rs (through the IP_3_-binding core) interact with phosphatidylinositol 4,5- bisphosphate (PIP_2_), which is dynamically associated with SOCE-MCS (Kang *et al*., 2019). We suggest that the extent to which IP_3_Rs tune SOCE in different cells is probably determined by the strength of Gq signaling and endogenous interactions between STIM1 and Orai1. The latter is likely to depend on the relative expression of STIM1 and Orai1 (Woo *et al*., 2020), the STIM isoforms expressed, expression of proteins that stabilize STIM1-Orai1 interactions (Darbellay *et al*., 2011; Rana *et al*., 2015; Rosado *et al*., 2016; Knapp *et al*., 2022), and the size and number of the MCS where STIM1 and Orai1 interact (Kang *et al*., 2019). The multifarious contributors to SOCE suggest that cells may differ in whether they express “spare capacity”. In neuronal cells, loss of IP_3_ (**Figure 4D**) or of the dominant IP_3_R isoform (IP_3_R1-shRNA; **Figures 1 and 2**) is sufficient to unveil the contribution of IP_3_R to SOCE, whereas HEK cells require loss of both IP_3_ and IP_3_R1 to unveil the contribution (**Figures 4H and 4I**). The persistence of SOCE in cells devoid of IP_3_Rs (**Figure 2 - figure supplement 3D and 3E**) (Prakriya and Lewis, 2001; Ma *et al*., 2002) probably arises from adaptive changes within the SOCE pathway. This does not detract from our conclusion that under physiological conditions, where receptors through IP_3_ initiate SOCE, IP_3_Rs actively regulate SOCE.

**Figure 8.**
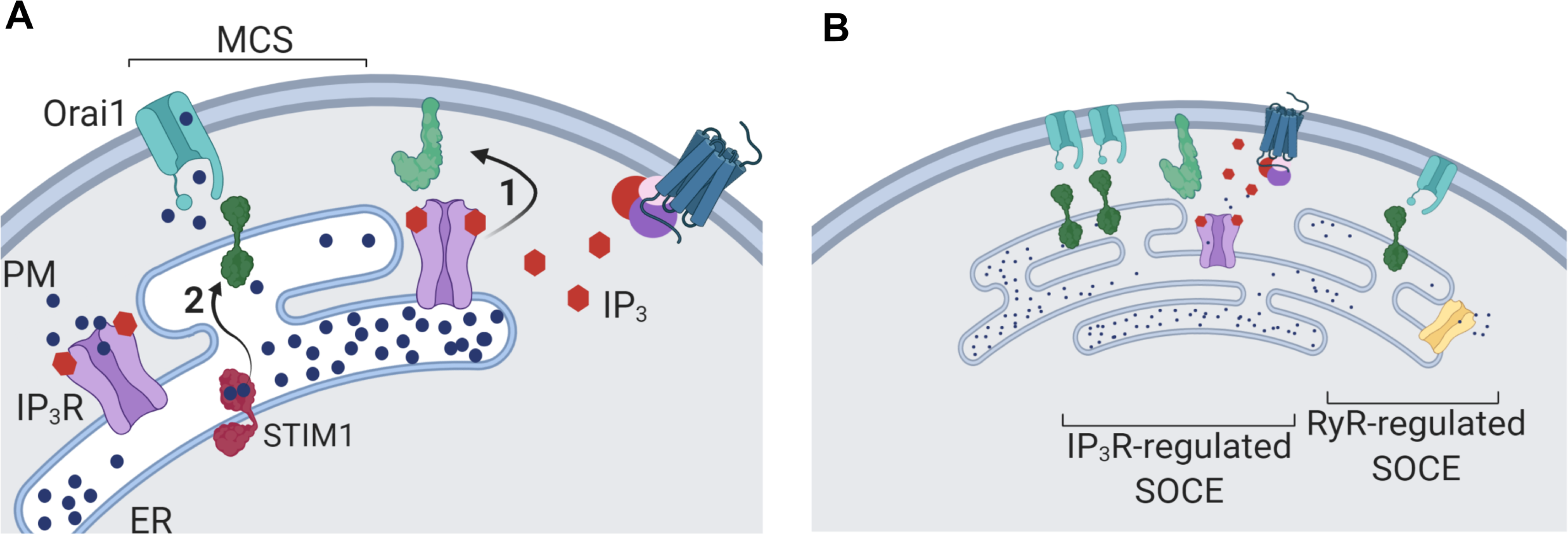
Dual Regulation of SOCE by IP_3_Rs. **(A)** SOCE is activated when loss of Ca^2+^ from the ER, usually mediated by opening of IP_3_Rs when they bind IP_3_, causes STIM to unfurl cytosolic domains (2). The exposed cytosolic domains of STIM1 reach across a narrow gap between the ER and PM at a MCS to interact with PIP_2_ and Orai1 in the PM. Binding of STIM1 to Orai1 causes pore opening, and SOCE then occurs through the open Orai1 channel. We show that IP_3_Rs when they bind IP_3_ also facilitate interactions between Orai1 and STIM, perhaps by stabilizing the MCS (1). Receptors that stimulate IP_3_ formation thereby promote both activation of STIM (by emptying Ca^2+^ stores) and independently promote interaction of active STIM1 with Orai1. **(B)** Other mechanisms, including ryanodine receptors (RyR), can also release Ca^2+^ from the ER. We suggest that convergent regulation of SOCE by IP_3_R with bound IP_3_ allows receptors that stimulate IP_3_ formation to selectively control SOCE.

The IP_3_Rs that initiate Ca^2+^ signals reside in ER immediately beneath the PM and alongside, but not within, the MCS where STIM1 accumulates after store depletion (Thillaiappan *et al*., 2017; **Figure 6A and 6B**). In migrating cells too, IP_3_Rs and STIM1 remain separated as they redistribute to the leading edge (Okeke *et al*., 2016). Furthermore, there is evidence that neither STIM1 nor STIM2 co-immmunoprecipitate with IP_3_R1 (Ahmad *et al*., 2022). We suggest, and consistent with evidence that SOCE in cells without IP_3_Rs can be restored by over-expressing E-Syt1 (**Figures 7A - 7C**), that ligand-bound IP_3_Rs probably facilitate SOCE by stabilizing the MCS wherein STIM1 and Orai1 interact, rather than by directly regulating either protein. This proposal provides an analogy with similar structural roles for IP_3_Rs in maintaining MCS between ER and mitochondria (Bartok *et al*., 2019) or lysosomes (Atakpa *et al*., 2018) (**Figure 8**). Since both contributions of IP_3_Rs to SOCE require IP_3_ binding (**Figures 3E and 3F**), each is ultimately controlled by receptors that stimulate IP_3_ formation (**Figures 4B and 4C**). Convergent regulation by IP_3_Rs at two steps in the SOCE pathway may ensure that receptor-regulated PLC activity provides the most effective stimulus for SOCE; more effective, for example, than ryanodine receptors, which are also expressed in neurons (**Figure 8B**). By opening IP_3_Rs parked alongside SOCE MCS (Thillaiappan *et al*., 2017; Ahmad *et al*., 2022), IP_3_ selectively releases Ca^2+^ from ER that is optimally placed to stimulate SOCE, and by facilitating Orai1- STIM1 interactions IP_3_ reinforces this local activation of SOCE (**Figure 8A and 8B**).

We conclude that IP_3_-regulated IP_3_Rs regulate SOCE by mediating Ca^2+^ release from the ER, thereby activating STIM1 and/or STIM2 (Ahmad *et al*., 2022) and, independent of their ability to release Ca^2+^, IP_3_Rs facilitate the interactions between STIM and Orai that activate SOCE. Dual regulation of SOCE by IP_3_ and IP_3_Rs allows robust control by cell-surface receptors and may reinforce local stimulation of Ca^2+^ entry.

## ACKNOWLEDGEMENTS

This research was supported by grants to G.H. from the Dept. of Biotechnology, Govt. of India (BT/PR6371/COE/34/19/2013) and NCBS-TIFR core support, to C.W.T. from the Wellcome Trust (101844) and Biotechnology and Biological Sciences Research Council (BB/T012986/1) and to D.I.Y from the NIH (NIDCR, DE014756). P.C is supported by a DST-INSPIRE fellowship (DST/INSPIRE Fellowship/2017/IF170360) and she received an Infosys-NCBS travel award to visit C.W.T.’s lab at Cambridge. We are grateful to Renjitha Gopurappilly (NCBS, TIFR) for the derivation of human neural precursor cells. We acknowledge use of the Central Imaging and Flow Cytometry Facility (CIFF), Stem Cell Culture Facility and Biosafety level-2 laboratory facility at NCBS, TIFR.

---

**Figure 1 - figure supplement 1.**
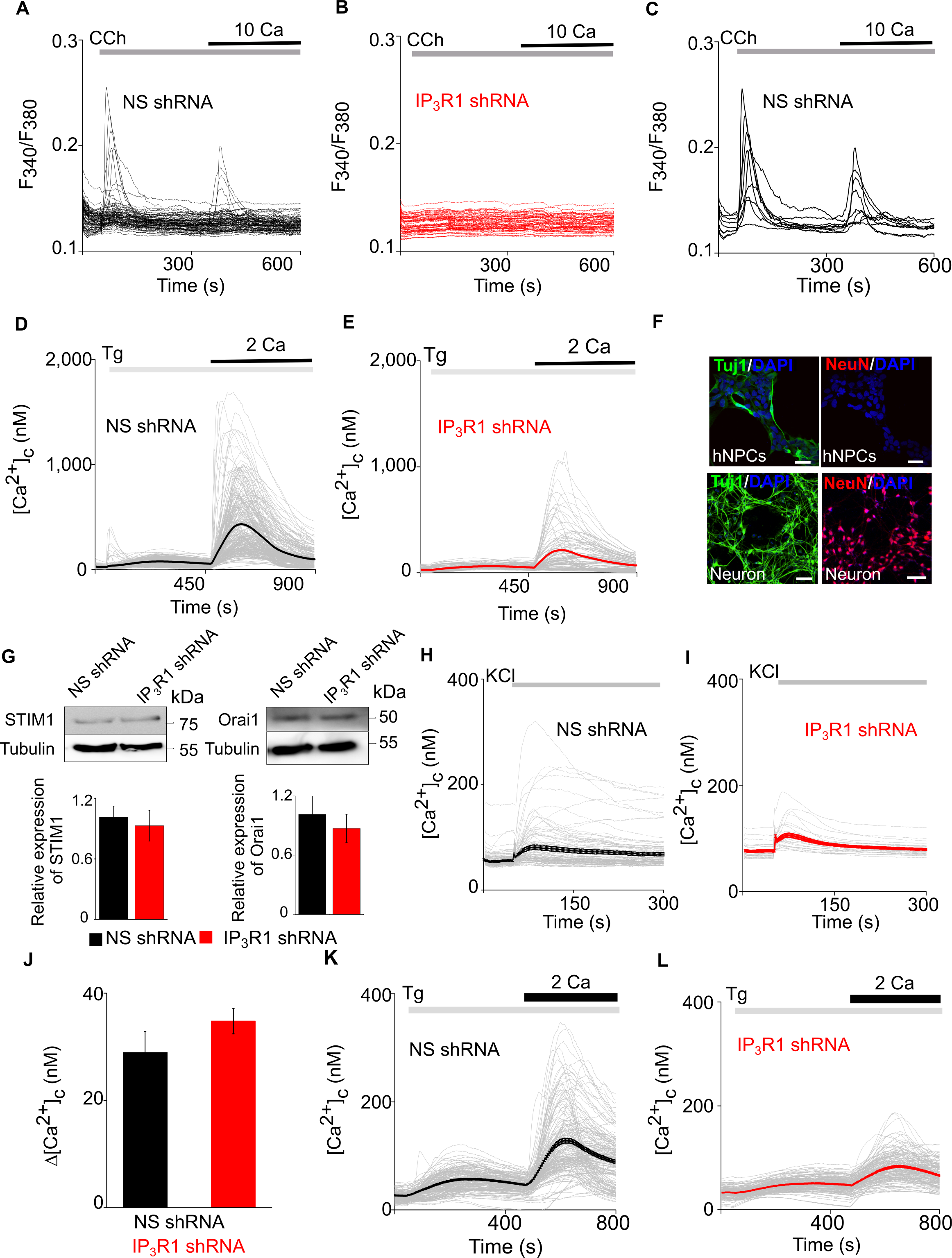
Loss of IP_3_R1 Attenuates SOCE In Neural Precursor Cells and Differentiated Neurons. (**A-C**) hNPCs expressing NS (**A, C**) or IP_3_R1-shRNA (**B**) were stimulated with carbachol (CCh, 100 µM) in Ca^2+^-free HBSS before restoring extracellular Ca^2+^ (10 mM). Each trace shows the Fura 2 fluorescence ratio (F_340_/F_380_) from a single cell (>50 cells from 3 experiments). Traces in (**C**) show only cells that responded to CCh. (**D, E**) hNPCs expressing shRNA were stimulated with thapsigargin (Tg, 10 µM) in Ca^2+^-free HBSS before restoring extracellular Ca^2+^ (2 mM). Traces are from individual cells (>180 cells from 3 experiments), with the mean response shown by a thick line. Summary results in **Figures 1D-1G**. (**F**) Confocal images of hNPCs and neurons spontaneously differentiated (10 days) from hNPCs stained for DAPI and neuronal markers: Tuj1 (βIII-tubulin) and NeuN (neuronal nuclear antigen). Scale bars, 50 μm. Only differentiated hNPCs express neuronal markers. (**G**) WB for STIM1 and Orai1 in lysates from hNPCs cells expressing non-silencing (NS) or IP_3_R1-shRNA. Summary results (mean ± s.d., n = 3) show relative expression of STIM1 and Orai1 relative to control shRNA (NS) cells. *P* > 0.01, Student’s *t*-test with unequal variances. (**H, I**) Spontaneously differentiated hNPCs (25 days) expressing NS (**H**) or IP_3_R1- shRNA (**I**) were depolarized by addition of KCl (75 mM). Traces show responses from single cells (>30 cells from 3 experiments) and mean response (thick line). (**J**) Summary results (mean ± s.e.m., 3 experiments) show peak increases in [Ca^2+^]_c_ (Δ[Ca^2+^]_c_) evoked by depolarization. No significant difference, Student’s *t*-test with unequal variances. (**K, L**) Spontaneously differentiated hNPCs expressing NS (**K**) or IP_3_R1-shRNA (**L**) were stimulated with Tg (10 µM) in Ca^2+^-free HBSS before restoring extracellular Ca^2+^ (2 mM). Traces from individual cells (>100 cells from 3 experiments) and the mean response (bold) are shown. Summary results in Figure 1H**-1J**. **Source data in Figure 1 - figure supplement 1 source data file.**

**Figure 2 - figure supplement 1.**
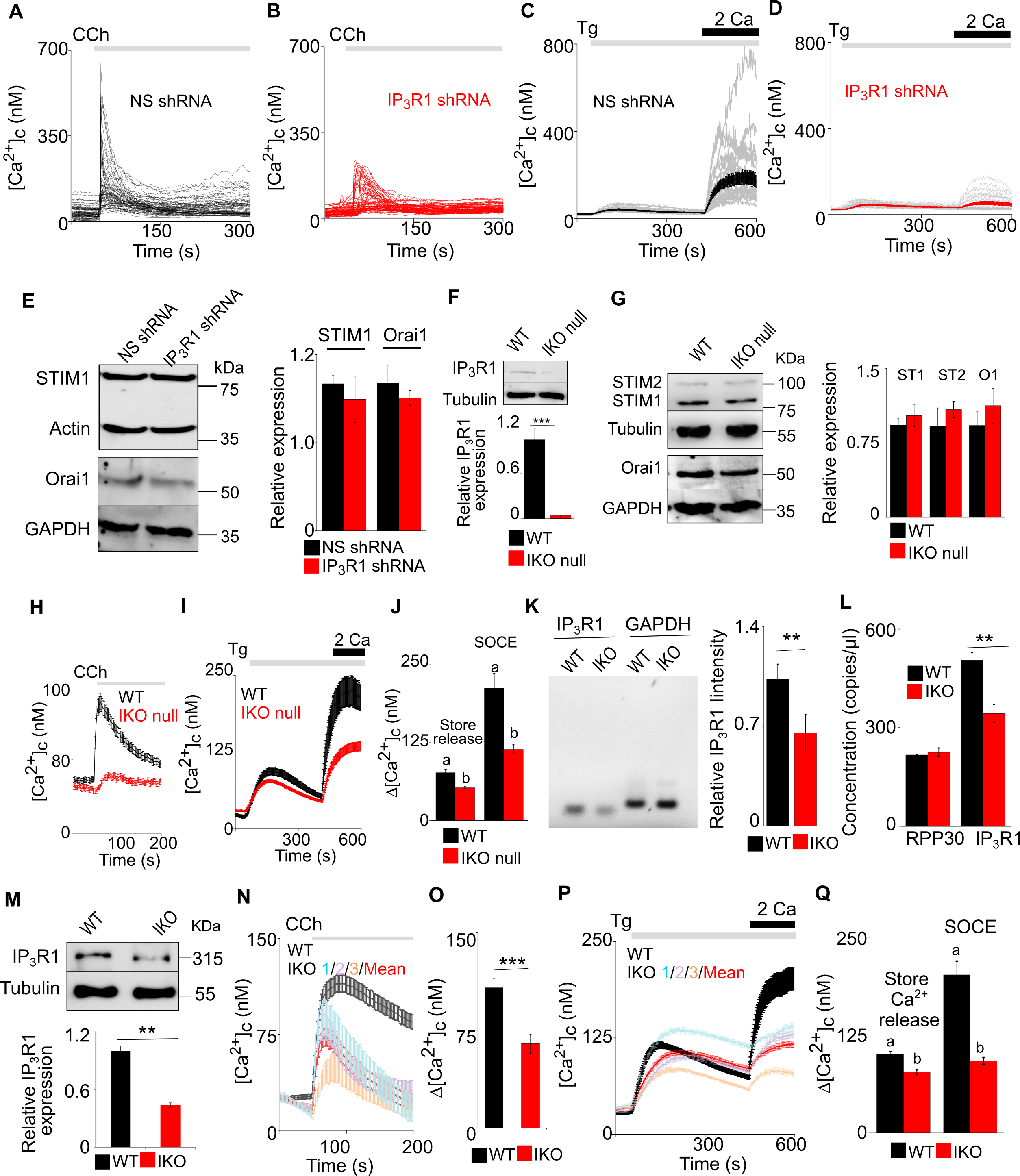
Reduced Expression of IP_3_R1 Using either shRNA or CRISPR/Cas9n Attenuates SOCE in SH-SY5Y Cells. (**A, B**) Ca^2+^ signals evoked by carbachol (CCh, 3 μM) in Ca^2+^-free HBSS in SH-SY5Y cells expressing NS (**A**) or IP_3_R1-shRNA (**B**). Traces show responses from single cells (100-200 cells from 3 experiments). (**C, D**) SH-SY5Y cells expressing NS (**C**) or IP_3_R1-shRNA (**D**) were treated with thapsigargin (Tg, 1 µM) in Ca^2+^-free HBSS before restoration of extracellular Ca^2+^ (2 mM). Traces show responses from single cells (50-100 cells from 3 experiments) and mean response (thick lines). Summary results in Figures 2C**-2G**. **(E)** Western blots (WB) for STIM1 and Orai1 from lysates of SH-SY5Y cells expressing non-silencing (NS) or IP_3_R1-shRNA. Summary results (mean ± s.d., n = 3) show STIM1 and Orai1 expression relative to NS-shRNA cells. *P* > 0.05, Student’s *t*- test with unequal variances. **(F)** Western blot for IP_3_R1 and tubulin from lysates from WT SH-SY5Y cells and for cells from a cell line in which the IP_3_R1 gene was targeted using CRISPR/Cas9 (IKO null). The blot is typical of 3 similar analyses. Summary results (mean ± s.d., n = 3) show IP_3_R1 expression relative to WT cells. P < 0.001, Student’s *t*-test with unequal variances. **(G)** Western blots (WB) for STIM1 (S1), STIM2 (S2) and Orai1 (O1) from lysates of WT and IKO null SH-SY5Y cells. Summary results (mean ± s.d., n = 3) show STIM1,2 and Orai1 expression relative to WT cells. P > 0.05, Student’s t-test with unequal variances. **(H)** Ca^2+^ signals evoked by carbachol (CCh, 1 μM) in WT and IKO null SH-SY5Y cells in Ca^2+^-free HBSS. **(I)** SH-SY5Y cells (WT or IKO null) were treated with thapsigargin (Tg, 1 µM) in Ca^2+^- free HBSS before restoring extracellular Ca^2+^ (2 mM). Mean ± s.e.m. from 3 experiments with WT cells (black, 110 cells), IKO null cells (red, 117 cells). **(J)** Summary results show the peak increases in [Ca^2+^]_c_ evoked by Tg (1 µM) in Ca^2+^- free HBSS (Δ[Ca^2+^]_c_) and after Ca^2+^ restoration (SOCE). Different letters indicate P < 0.001, Mann-Whitney U-test. **(K)** PCRs for IP_3_R1 and GAPDH (control gene) of genomic DNA isolated from SH- SY5Y cells. Results (mean ± s.e.m. from 3 independent analyses of a single cell line) are shown for wild type (WT) cells and for cells in which the IP_3_R1 gene was targeted using CRISPR/Cas9n (IKO). Expression of IP_3_R1 has been normalized to expression of GAPDH. \*\**P* < 0.01, Student’s *t*-test with unequal variances. **(L)** Droplet digital PCR amplification was used to determine gene copy number for IP_3_R1 and a control gene RPP30 for WT and IKO SH-SY5Y cells. \*\**P* < 0.05, Student’s *t*-test with unequal variances. **(M)** Western blot for IP_3_R1 and tubulin from lysates from WT SH-SY5Y cells and for cells from a cell line in which the IP_3_R1 gene was targeted using CRISPR/Cas9n (IKO). The blot is typical of 3 similar analyses. Summary results (mean ± s.d., n = 3) show IP_3_R1 expression relative to WT cells. **P < 0.01, Student’s *t*-test with unequal variances. **(N)** Ca^2+^ signals evoked by carbachol (CCh, 1 μM) in WT and IKO SH-SY5Y cells in Ca^2+^-free HBSS. Results show (mean ± s.e.m.) for each of three independently edited cell lines (IKO1, 2 and 3), each with at least 30 cells; and the overall mean ± s.e.m. (red line). Results from a total of 89 WT cells and 130 IKO cells are shown. **(O)** Summary results (mean ± s.e.m. from 3 experiments with 80-100 cells) show peak changes in [Ca^2+^]_c_ (Δ[Ca^2+^]_c_) evoked by CCh. \*\*\**P* < 0.001, Mann-Whitney U-test. **(P)** SH-SY5Y cells (WT or IKO) were treated with thapsigargin (Tg, 1 µM) in Ca^2+^-free HBSS before restoring extracellular Ca^2+^ (2 mM). Mean ± s.e.m. from 3 experiments with WT cells (black, 85 cells), 3 independently edited cell lines (IKO 1, 2 and 3; <u>at least</u> 30 cells for each) and the pooled results from all edited cells (97; red). **(Q)** Summary results show the peak increases in [Ca^2+^]_c_ evoked by Tg (1 µM) in Ca^2+^- free HBSS (Δ[Ca^2+^]_c_) and after Ca^2+^ restoration (SOCE). Different letters indicate P < 0.001, Mann-Whitney U-test. **Source data in Figure 2 - figure supplement 1 source data file.**

**Figure 2 - figure supplement 2.**
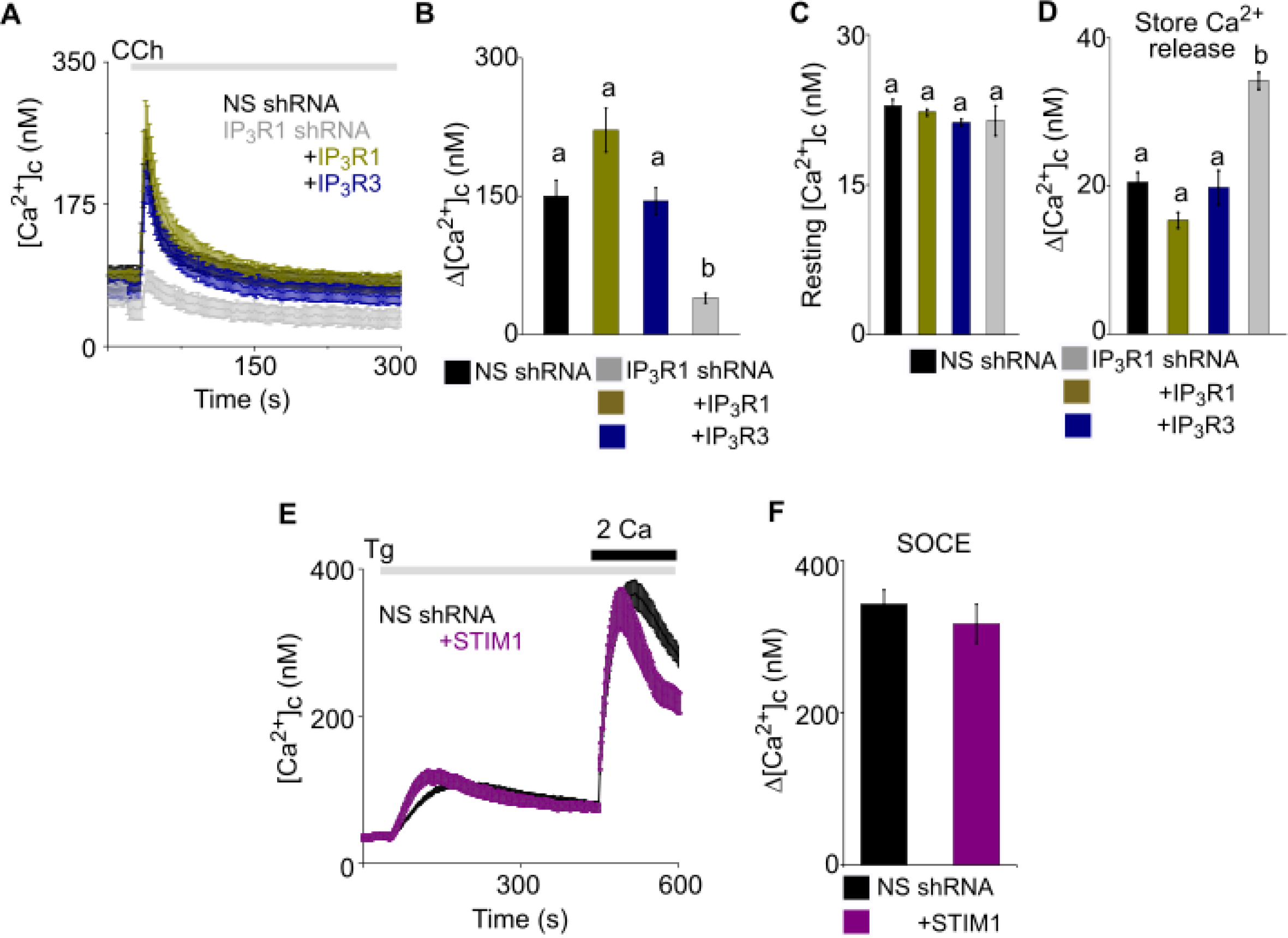
Attenuated SOCE In SH-SY5Y Cells Lacking IP_3_R1 is Rescued by Expression of IP_3_R1, IP_3_R3 or STIM1. **(A)** Ca^2+^ signals evoked by carbachol (CCh, 3 μM) in SH-SY5Y cells expressing NS or IP_3_R1-shRNA, and the latter with IP_3_R1 or IP_3_R3. Traces show mean ± s.e.m. from >3 experiments with 50-70 cells. (**B-D**) Summary results show peak Ca^2+^ signals (Δ[Ca^2+^]_c_) evoked by CCh (**B**), resting [Ca^2+^]_c_ (**C**) and the peak Ca^2+^ signal evoked by Tg (10 µM) in Ca^2+^-free HBSS (Δ[Ca^2+^]_c_, a reporter of ER Ca^2+^ content) (**D**). Different letter codes indicate significantly different values, *P* < 0.001, one-way ANOVA and pair-wise Tukey’s test. (**E**) Effects of expressing mCh-STIM1 in SH-SY5Y cells expressing NS shRNA on the Ca^2+^ signals evoked by Tg (1 µM) in Ca^2+^-free HBSS and then after restoration of extracellular Ca^2+^ (2 mM). Traces show mean ± s.e.m. from 3 experiments with 30- 110 cells. (**F**) Summary results show the peak Δ[Ca^2+^]_c_ evoked by restoring extracellular Ca^2+^ after Tg. Mean ± s.e.m. *P* > 0.05, Mann-Whitney U-test. **Source data in Figure 2 - figure supplement 2 source data file.**

**Figure 2 - figure supplement 3.**
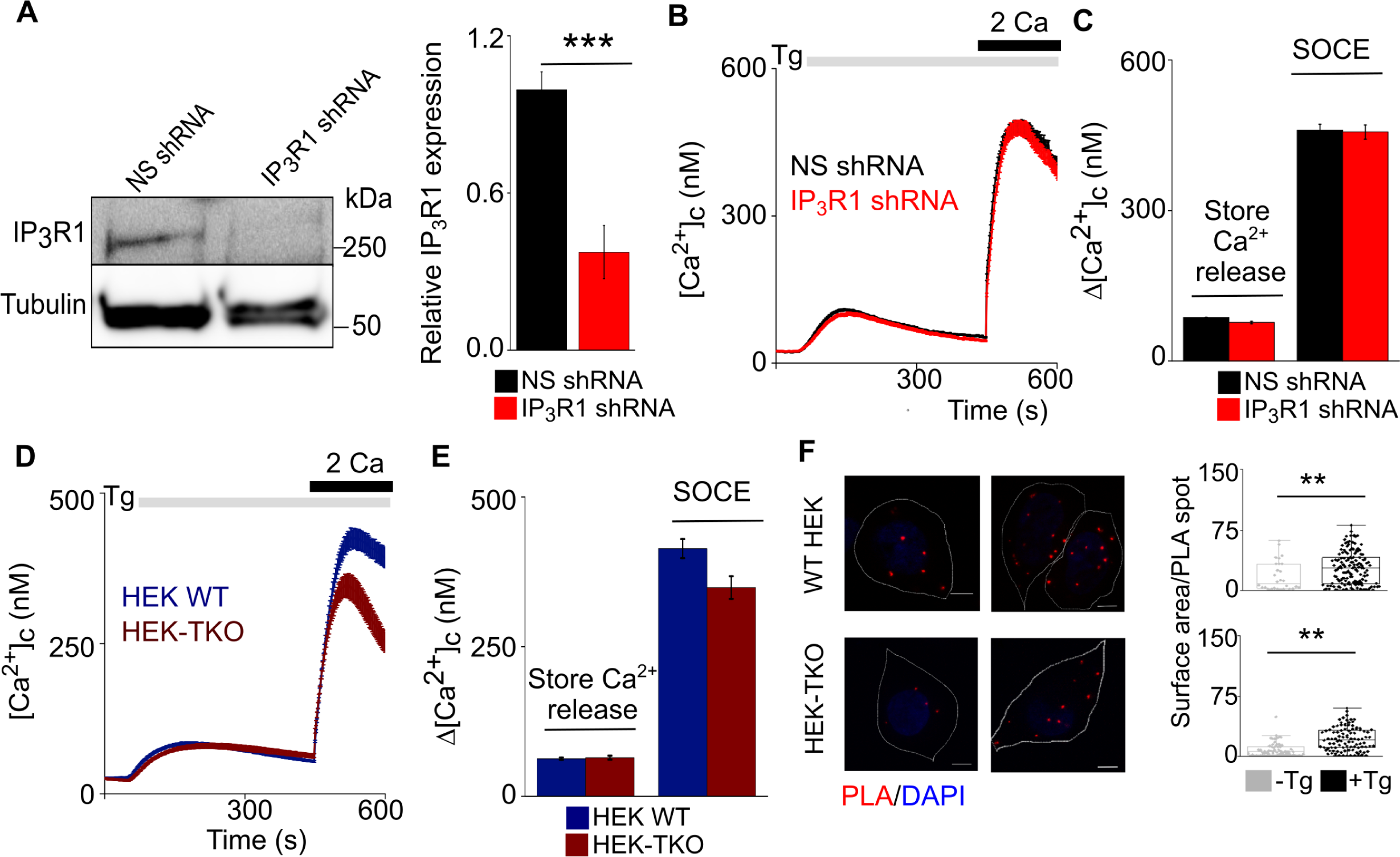
Loss of IP_3_R1 Does Not affect SOCE in HEK Cells. **(A)** WB for IP_3_R1 in HEK cells expressing non-silencing (NS) or IP_3_R1-shRNA. Summary results (mean ± s.d., n = 3) show IP_3_R1 expression relative to tubulin. \*\*\**P* < 0.001, Student’s *t*-test with unequal variances. **(B)** HEK cells expressing NS or IP_3_R1-shRNA were treated with thapsigargin (Tg, 1 µM) in Ca^2+^-free HBSS before restoration of extracellular Ca^2+^ (2 mM). Traces show mean response from >100 cells from 3 experiments. **(C)** Summary results show the peak increases in [Ca^2+^]_c_ evoked by Tg (1 µM) in Ca^2+^- free HBSS (Δ[Ca^2+^]_c_, a reporter of ER Ca^2+^ content) and after Ca^2+^ restoration (SOCE). **(D)** Wild type HEK cells (HEK WT) and cells lacking all IP_3_Rs (HEK-TKO) were treated with thapsigargin (Tg, 1 µM) in Ca^2+^-free HBSS before restoration of extracellular Ca^2+^ (2 mM). Traces show mean ± s.e.m. from 3 experiments with 80-120 cells. **(E)** Summary results show the peak Δ[Ca^2+^]_c_ evoked by restoring extracellular Ca^2+^ after Tg. Mean ± s.e.m. *P* > 0.01, Mann-Whitney U-test. **(F)** Proximity ligation assay (PLA) analyses of interactions between STIM1 and Orai1 in wild type HEK (WT) and TKO cells. Confocal images are shown for control cells (- Tg) or after10 min treatment with thapsigargin (+Tg, 1 µM) in Ca^2+^-free HBSS. PLA reaction product is red, and nuclei are stained with DAPI (blue). Scale bars, 5 µm. Summary results show the surface area of the PLA spots for ∼20 cells from 2 independent analyses. Individual values, median (bar) and 25th and 75th percentiles (box). **P < 0.01, Student’s t-test with unequal variances. **Source data in Figure 2 - figure supplement 3 source data file.**

**Figure 3 - figure supplement 1.**
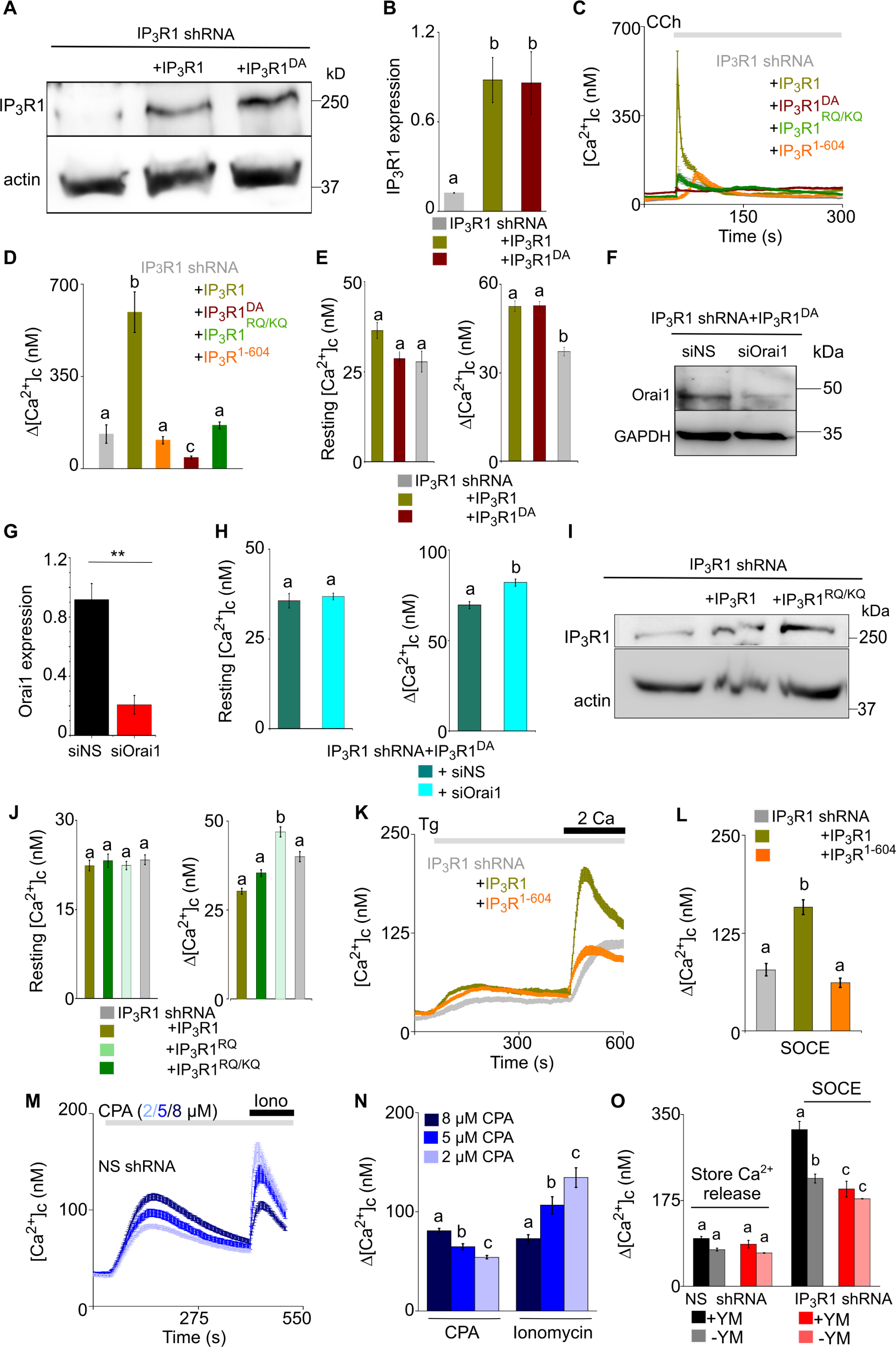
Attenuated SOCE In SH-SY5Y Cells Lacking IP_3_R1 is Rescued by Expression of Pore-Dead IP_3_R1 with a Functional IP_3_-Binding Site. **(A)** WB with IP_3_R1 antibody showing expression of IP_3_R1 and IP_3_R1^DA^ in SH-SY5Y cells expressing IP_3_R1-shRNA. **(B)** Summary (mean ± s.e.m, n = 3) shows IP_3_R1 expression normalized to actin. *P* < 0.001, one-way ANOVA with pair-wise Tukey’s test. Here, and in most subsequent panels, different letters indicate groups that are statistically different, with the test defined for each panel. **(C)** Ca^2+^ signals evoked by carbachol (CCh 3 μM) in SH-SY5Y cells expressing IP_3_R1- shRNA alone or with IP_3_R1, IP_3_R1^DA^, IP_3_R1^RQ/KQ^ or IP_3_R1^1-604^. Traces show mean ± s.e.m. from 3 experiments. **(D)** Summary results (mean ± s.e.m) show peak changes in [Ca^2+^]_c_ evoked by CCh (Δ[Ca^2+^]_c_). *P* < 0.001, one-way ANOVA with pair-wise Tukey’s test. **(E)** Resting [Ca^2+^]_c_ (left) and thapsigargin (Tg)-evoked Ca^2+^ release (1 µM, Δ[Ca^2+^]_c_) (right) in cells treated as indicated. *P* < 0.001, one-way ANOVA with pair-wise Tukey’s test. **(F)** WB showing Orai1 expression in cells expressing IP_3_R1-shRNA and IP_3_R1^DA^, and then treated with NS or Orai1-siRNA. **(G)** Summary results (normalized to GAPDH expression, mean ± s.d, *n* =3). *\*\*P* < 0.01, Student’s *t*-test with unequal variances. **(H)** Resting [Ca^2+^]_c_ (left) and Tg-evoked Ca^2+^ release (right) in cells expressing IP_3_R1- shRNA and IP_3_R1^DA^, and then treated with NS or Orai1-siRNA. *P* < 0.05, Mann- Whitney U-test. **(I)** WB for IP_3_R1 in cells expressing IP_3_R1-shRNA alone or with IP_3_R1 or IP_3_R1^RQ/KQ^. Results typical of 2 independent experiments. **(J)** Resting [Ca^2+^]_c_ (left) and Ca^2+^ release evoked by Tg (right) in cells expressing IP_3_R1-shRNA alone or with the indicated IP_3_R1. Mean ± s.e.m., *n* = 3 experiments. *P* < 0.001, one-way ANOVA with pair-wise Tukey’s test. **(K)** Changes in [Ca^2+^]_c_ evoked by Tg (10 µM) in Ca^2+^-free HBSS and then after restoration of extracellular Ca^2+^ (2 mM) in cells expressing IP_3_R1-shRNA alone or with IP_3_R1 or IP_3_R1^1-604^. Mean ± s.e.m. from 3 experiments with >100 cells. **(L)** Summary results show peak increases in [Ca^2+^]_c_ evoked by Ca^2+^ restoration in the indicated cells. *P* < 0.001, one-way ANOVA with pair-wise Tukey’s test. **(M)** SH-SY5Y cells expressing NS-shRNA were stimulated with the indicated concentrations of CPA in Ca^2+^-free HBSS to partially deplete ER Ca^2+^ stores and then with ionomycin (Iono, 1 µM) to release all remaining Ca^2+^. Results show mean ± s.e.m. for >30 cells from 3 experiments. **(N)** Summary results (mean ± s.e.m.) show Δ[Ca^2+^]_c_ evoked by CPA or ionomycin. *P* < 0.001, one-way ANOVA with pair-wise Tukey’s test. The results confirm that the lower concentrations of CPA partially deplete the ER of Ca^2+^. **(O)** Summary results (mean ± s.e.m, 80-100 cells from 3 experiments) show the peak increases in [Ca^2+^]_c_ (Δ[Ca^2+^]_c_) after adding Tg (store Ca^2+^ release) and after restoring extracellular Ca^2+^ (SOCE) with or without YM-254890 treatment. Different letter codes indicate significantly different values within the Ca^2+^ release or SOCE groups, *P* < 0.001, one-way ANOVA and pair-wise Tukey’s test. **Source data in Figure 3 - figure supplement 1 source data file.**

**Figure 4 - figure supplement 1.**
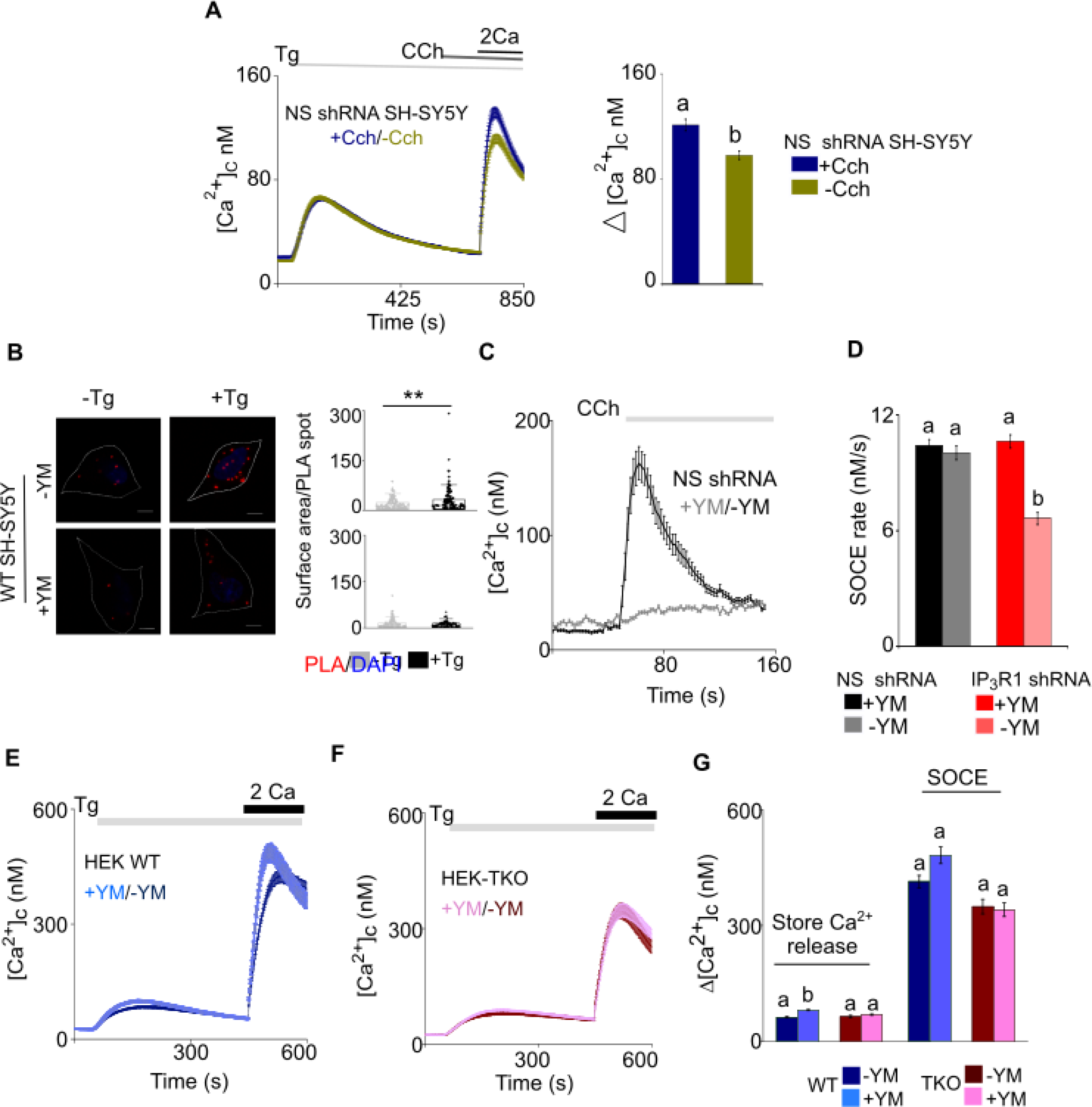
Effects of Generating IP_3_ and Inhibiting Gq on Ca^2+^ Signals and STIM1-Orai1 interactions in SH-SY5Y and HEK Cells. **(A)** SOCE is enhanced in SH-SY5Y cells expressing control-shRNA after ER-Ca^2^ depletion (thapsigargin, 2 µM) followed by carbachol (CCh, 1 µM) stimulated IP_3_ formation. Traces (mean ± s.e.m of ∼270 cells each from 5 experiments) show responses with and without the CCh addition. Summary results show the peak increases in [Ca^2+^]_c_ (Δ[Ca^2+^]_c_) for SOCE after restoring extracellular Ca^2+^. Different letters indicate significantly different values of P < 0.001 by Mann- Whitney U-test. **(B)** PLA analyses of interactions between STIM1 and Orai1 in wild type (WT) SH- SY5Y cells untreated (-YM) or treated (+YM, 1μm) with Gq inhibitor YM-254890. Confocal images are shown for control cells (-Tg) or after treatment with thapsigargin (+Tg, 1 µM) in Ca^2+^-free HBSS. PLA reaction product is red, and nuclei are stained with DAPI (blue). Scale bars, 5 µm. Summary results show the surface area of the PLA spots for ∼20 cells from 2 independent analyses. Individual values, median (bar) and 25th and 75th percentiles (box). **P < 0.01, Student’s t-test with unequal variances. **(C)** HEK cells expressing NS shRNA and treated with the Gq inhibitor, YM-254890 (YM, 1 μM, 5 min) were stimulated with carbachol (CCh, 100 μM). Traces show mean ± s.e.m. from 3 experiments with 30-40 cells. The results confirm that treatment with YM-254890 effectively uncouples CCh from IP_3_-evoked Ca^2+^ signals. **(D)** From results shown in Figures 4G and 4H, rates of thapsigargin-evoked Ca^2+^ entry were determined. Summary shows mean ± s.e.m, from 100-120 cells. *P* < 0.001, Mann-Whitney U-test. (**E-F**) Changes in [Ca^2+^]_c_ evoked by thapsigargin (Tg, 1 μM) in Ca^2+^-free HBSS and then after restoration of extracellular Ca^2+^ (2 mM) in WT (**C**) and TKO (**D**) HEK cells after treatment with YM-254890 (YM, 1 μM, 5 min). Traces show mean ± s.e.m. from 3 experiments with 100-120 cells. (**G**) Summary results show peak increase in [Ca^2+^]_c_ (Δ[Ca^2+^]_c_) evoked by addition of thapsigargin and then after restoring extracellular Ca^2+^ (SOCE). Mean ± s.e.m. Different letter codes indicate significantly different values, *P* < 0.001, Mann- Whitney U-test. **Source data in Figure 4 - figure supplement 1 source data file.**

**Figure 5 - figure supplement 1.**
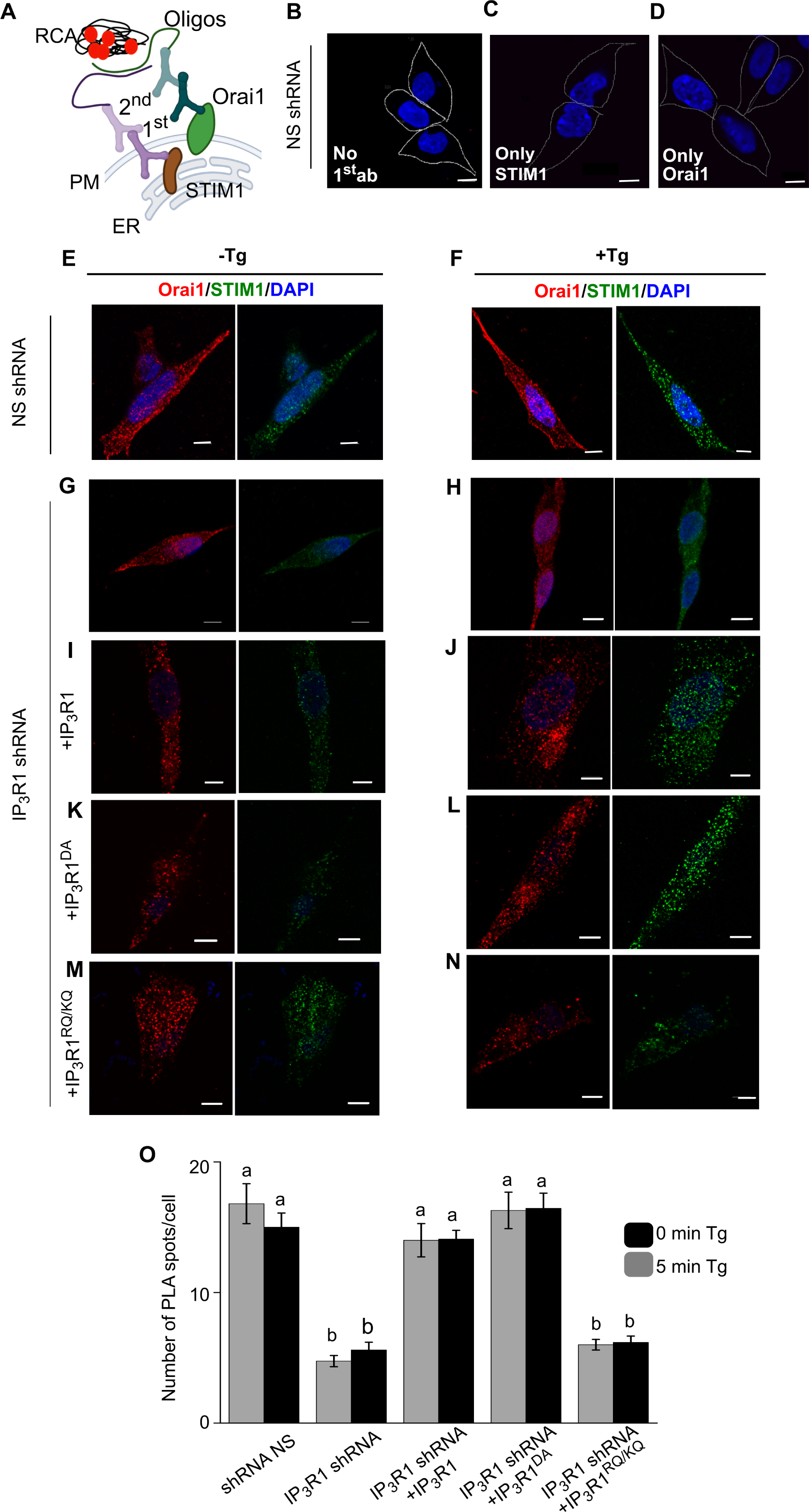
Validation of PLA Measurements of Orai1-STIM1 Interactions. (**A**) PLA using antibodies to Orai1 and STIM1 to establish whether they are associated. Complementary oligonucleotides conjugated to a secondary antibody (2^nd^) targeting their respective primary antibodies (1^st^) hybridize when they are close to each other, allowing rolling circle amplification (RCA) and production of a red fluorescent product. The latter can be quantified from the surface area of the fluorescent spots or their intensity. We obtained indistinguishable results with both methods of quantification. We therefore show only results obtained from surface-area measurements. (**B-D**) Confocal images from PLA assays of SH-SY5Y cells expressing NS-shRNA performed with no primary antibody (**B**), or with primary antibody for only STIM1 (**C**) or only Orai1 (**D**). Scale bars, 5 µm. (**E, F**) Confocal images of SH-SY5Y cells expressing NS-shRNA with (**F**) or without thapsigargin treatment (**E**) (Tg, 1 µM, in Ca^2+^-free HBSS). Cells are stained with DAPI and immunostained for Orai1 or STIM1. Scale bars, 5 µm. **(G-N)** Similar analyses of cells expressing IP_3_R1-shRNA alone (**G, H**), or with IP_3_R1 (**I, J**), IP_3_R1^DA^ (**K, L**) or IP_3_R1^RQ/KQ^ (**M, N**). Results (**E-N**) are typical of 2 experiments. **(O)** Number of PLA spots per cell in the indicated genotypes with (5 min) or without (0 min) thapsigargin treatment (Tg, 1 µM in Ca^2+^-free HBSS) in the indicated genotypes. Different letter indicates significantly different values, P < 0.001 one-way ANOVA and pair-wise Tukey’s test. The PLA results are shown in Figure 5.

**Figure 6 - figure supplement 1.**
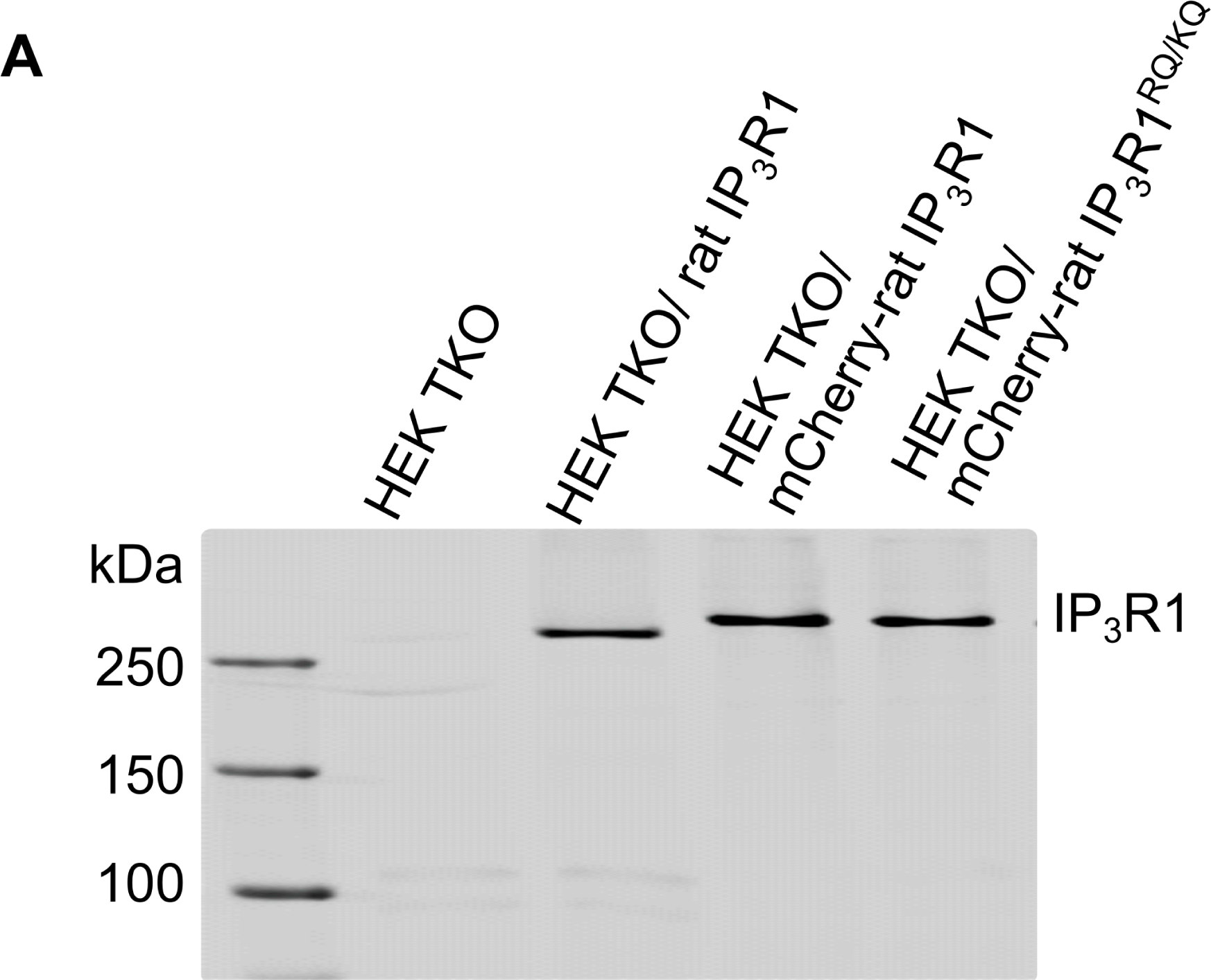
Validation of fluorescent-tagged rat IP_3_R1 constructs. **(A)** Western blot image showing the expression of IP_3_R1 in the indicated genotypes Results with these constructs are shown in Figure 6. **Source data in Figure 6 – figure supplement 1.**

## RESOURCE AVAILABILITY

### Lead Contact

All requests for resources and reagents should be directed to the lead contact, Dr. Gaiti Hasan.

### Materials Availability

Constructs and cell lines are available upon request. MTA required for cell lines.

### Data and Code Availability

This study did not generate any computer code. The data supporting the findings of this study are available within the manuscript. All other data supporting the findings of this study are available in source data file of respective figures.

## METHOD DETAILS

### Culture of Human Neural Precursor Cells

Human neural precursor cells (hNPCS) were derived from a human embryonic stem cell (hESC) line, H9/WA09 (RRID: CVCL_9773), using a protocol that inhibits dual SMAD signalling and stimulates Wnt signaling (Reinhardt *et al*., 2013) as described previously (Gopurappilly *et al*., 2018, 2019). hNPCs were grown as adherent dispersed cells on growth factor-reduced Matrigel (0.5%, Corning, Cat#356230) in hNPC maintenance medium (NMM) at 37°C in humidified air with 5% CO_2_. NMM comprised a 1:1 mixture of Dulbecco’s Modified Eagle Medium with Nutrient Mixture F-12 (DMEM/F-12, Invitrogen, Cat#10565018) and Neurobasal medium (ThermoFisher, Cat#21103049), supplemented with GlutaMAX (0.5x, ThermoFisher, Cat#35050061), N2 (1:200, ThermoFisher, 17502048), B27 without vitamin A (1:100, ThermoFisher, Cat#12587010), Antibiotic-Antimycotic (ThermoFisher, Cat#15240112), CHIR99021 (3 μM, STEMCELL Technologies, Cat#72052), purmorphamine (0.5 mM, STEMCELL Technologies, Cat#72202) and ascorbic acid (150 μM, Sigma, Cat#A92902). Doubling time was ∼24 hr. Cells were passaged every 4-5 days by treatment with StemPro Accutase (ThermoFisher, Cat#A1110501), stored in liquid nitrogen, and thawed as required. Cells were confirmed to be mycoplasma- free by monthly screening (MycoAlert, Lonza, Cat#LT07-318). hNPCs between passages 16-19 were used.

All experiments performed with hESC lines were approved by the Institutional Committee for Stem Cell Research, registered under the National Apex Committee for Stem Cell Research and Therapy, Indian Council of Medical Research, Ministry of Health, New Delhi.

### Stable Knockdown of IP_3_R1

An UltramiR lentiviral inducible shRNA-mir based on the shERWOOD algorithm (Auyeung *et al*., 2013; Knott *et al*., 2014) was used to inhibit IP_3_R1 expression. The all-in-one pZIP vector, which allows puromycin-selection and doxycycline-induced expression of both shRNA-mir and Zs-Green for visualization, was from TransOMIC Technologies (Huntsville, AL). Lentiviral pZIP transfer vectors encoding non-silencing shRNA (NS, NT#3-TTGGATGGGAAGTTCACCCCG) or IP_3_R1-targeting shRNA (ULTRA3316782- TTTCTTGATCACTTCCACCAG) were packaged as lentiviral particles using packaging (pCMV- dR8.2 dpvr, Addgene, plasmid #8455) and envelope vectors (pCMV-VSV-G, Addgene, plasmid #8454) by transfection of HEK293T cells (referred as HEK, ATCC, Cat# CRL-3216). Viral particles were collected and processed and hNPCs (passage 9) or SH-SY5Y cells were transduced (multiplicity of infection, MOI = 10) using Lipofectamine LTX with PLUS reagent (ThermoFisher, Cat#15338100). Cells were maintained in media containing doxycycline (2 μg/ml, Sigma, Cat# D3072) to induce shRNA expression, and puromycin to select transduced cells (1 μg/ml for hNPCs; 3 μg/ml for SH-SY5Y cells; Sigma, Cat# P9620). Cells were passaged 4-5 times after lentiviral transduction to select for stable expression of shRNAs.

### Derivation of Neurons From hNPCs

Neurons were differentiated from hNPCs stably transduced with shRNA. hNPCs were seeded at 50-60% confluence in NMM on coverslips coated with poly-d-lysine (0.2 mg/ml, Sigma, Cat#P7280). After 1-2 days, the medium was replaced with neuronal differentiation medium, which comprised a 1:1 mixture of DMEM/F-12 with Neurobasal supplemented with B27 (1:100), N2 (1:200), GlutaMAX (0.5x) and Antibiotic-Antimycotic solution. Medium was replaced on alternate days. Neurons were used after 15-20 days.

### Culture and Transfection of SH-SY5Y Cells

SH-SY5Y cells (ATCC, USA, Cat# CRL-2266) were grown on culture dishes in DMEM/F-12 with 10% fetal bovine serum (Sigma, Cat# F4135) at 37 °C in humidified air with 5% CO_2_. Cells were passaged every 3-4 days using TrypLE Express (ThermoFisher, Cat# 12605036) and confirmed to be free of mycoplasma. Cells expressing shRNA were transiently transfected using TransIT-LT1 reagent (Mirus, Cat# MIR-2300) in Opti-MEM (ThermoFisher, Cat# 31985062). Plasmids (250 ng) and/or siRNA (200 ng) in transfection reagent (1 µg/2.5 µl) were added to cells grown to 50% confluence on glass coverslips attached to an imaging dish. Cells were used after 48 hrs. The siRNAs used were to human Orai1 (100 nM, Dharmacon, Cat# L- 014998-00-0005) or non-silencing (NS, Dharmacon, Cat# D-495 001810-10-05), to human STIM1 (Santa Cruz Biotechnology, Cat# sc-76589) or NS (Santa Cruz Biotechnology, Cat# sc-37007). The expression plasmids were IP_3_R1 (rat type 1 IP_3_R1 in pcDNA3.2/V5DEST vector) (Dellis *et al*., 2008), rat IP_3_R1^DA^ (D2550 replaced by A in pcDNA3.2 vector) (Dellis *et al*., 2008), rat IP_3_R1^RQ^ (R568 replaced by Q of type 1 IP_3_R in pCDNA3.2/V5DEST vector) (Dellis *et al*., 2008), rat IP_3_R1^RQ/KQ^ (R568 and K569 replaced by Q of type 1 IP_3_R in pCDNA3.2/V5DEST vector), rat IP_3_R1^1-604^ (residues 1-604 of IP_3_R with N-terminal GST tag in pCDNA3.2/V5DEST vector) (Dellis *et al*., 2008), rat IP_3_R3 (rat type 3 IP_3_R in pcDNA3.2/V5DEST vector) (Saleem *et al*., 2013), human mCherry-STIM1 (N terminal mCherry tagged human STIM1 in pENTR1a vector) (Nunes-Hasler *et al*., 2017) and human extended synaptotagmin 1 (E-Syt1), a kind gift from Dr S. Muallem, NIDCR, USA (Maléth *et al*., 2014).

### CRISPR/Cas9 and Cas9n Editing of SH-SY5Y Cells

To allow either CRISPR/Cas9 or Cas9n-mediated disruption of IP_3_R1 expression, we used a published method to clone gRNAs into the backbone vector (pSpCas9n(BB)- 2A-Puro PX462 V2.0, Addgene, Cat#62987) (Ran *et al*., 2013). Forward and reverse sgRNA oligonucleotides (100 µM) were annealed and ligated using T4 DNA ligase by incubation (10 µl, 37°C, 30 min) before slow cooling to 20°C. Plasmids encoding Cas9n were digested with *BbsI-HF* (37°C, 12 hr), gel-purified (NucleoSpin Gel and PCR Clean-up kit from Takara) and the purified fragment was stored at -20°C. A mixture (final volume 20 µl) of gRNA duplex (1 µl, 0.5 µM), digested px459 (for IKO null) or pX462 vector (for IKO) (30 ng), 10× T4 DNA ligase buffer (2 µl) and T4 DNA ligase (1 µl) was incubated (20°C, 1 hr). After transformation of DH5-α competent *E. coli* with the ligation mixture, plasmids encoding Cas9 or Cas9n and the sgRNAs were extracted, and the coding sequences were confirmed (Ran *et al*., 2013). The plasmid (2 µg) was then transfected into SH-SY5Y cells (50-60% confluent) in a 6-well plate using TransIT LT-1 reagent (Mirus Bio, Cat# MIR-2300). After 48 hr, puromycin (3 µg/ml, 72 hr) was added to kill non-transfected cells. IKO colonies were propagated and screened for Ca^2+^ signals evoked by carbachol and for the presence of the IP_3_R gene by genomic DNA PCR and droplet digital PCR using primers close to the region targeted by the gRNAs (Miotke *et al*., 2014). Three independently derived IKO lines, each with one residual IP_3_R1 gene, were used for analyses of Ca^2+^ signaling (see **Figure S2N-S2Q**). For one of the cell lines (IKO 2), disruption of one copy of the IP_3_R1 gene was confirmed by genomic PCR, droplet digital PCR and Western blotting (**Figures S2K-S2M**). For the IKO null line, single cell selection was done in a 96 well plate setup followed by screening for carbachol-evoked Ca^2+^ signals from multiple clones. A single clone was selected (**Figure S2H**) and a western blot performed to confirm absence of IP_3_R1 expression (**Figure S2F**). All the oligonucleotide sequences are described in **supplementary file 1.**

### Plasmid construction

Mutagenesis and all DNA modifications were carried out using *Q5* Hot Start high-fidelity 2X Master Mix (New England BioLabs, Cat# M0494L) using the recommendations of the manufacturer. Primers used in this study (details given in **supplementary file 1**) were synthesized by Integrated DNA Technologies (IDT). Mutations in the Ligand binding domain (R568Q and K569Q) of IP_3_R1 were generated on the rat mCherry-IP_3_R1 cDNA in pDNA3.1 Mutations in all the constructs were confirmed by sequencing.

### Ca^2+^ Imaging

Methods for single-cell Ca^2+^ imaging were described previously (Gopurappilly *et al*., 2019). Briefly, cells grown as a monolayer (∼70% confluence) on homemade coverslip-bottomed dishes were washed and loaded with Fura 2 by incubation with Fura 2 AM (4 μM, 45 min, 37°C, ThermoFisher, Cat# F1221), washed and imaged at room temperature in HEPES-buffered saline solution (HBSS). HBSS comprised: 20 mM HEPES, 137 mM NaCl, 5 mM KCl, 2 mM MgCl_2_, 2 mM CaCl_2_, 10 mM glucose, pH 7.3. CaCl_2_ was omitted from Ca^2+^-free HBSS. Treatments with carbachol (CCh, Sigma, Cat# C4382), thapsigargin (Tg, ThermoFisher, Cat# 7458), cyclopiazonic acid (CPA, Sigma Cat# C1530) or high-K^+^ HBSS (HBSS supplemented with 75 mM KCl) are described in legends.

Responses were recorded at 2-s intervals using an Olympus IX81-ZDC2 Focus Drift-Compensating Inverted Microscope with 60× oil immersion objective (numerical aperture, NA = 1.35) with excitation at 340 nm and 380 nm. Emitted light (505 nm) was collected with an Andor iXON 897E EMCCD camera and AndoriQ 2.4.2 imaging software (RRID: SCR_014461). Maximal (R_max_) and minimal (R_min_) fluorescence ratios were determined by addition of ionomycin (10 μM, Sigma, Cat# 407953) in HBSS containing 10 mM CaCl_2_ or by addition of Ca^2+^-free HBSS containing BAPTA (10 mM, Sigma, Cat# 196418) and Triton X100 (0.1%). Background-corrected fluorescence recorded from regions of interest ROI) drawn to include an entire cell was used to determine mean fluorescence ratios (R = F_340_/F_380_) (ImageJ), and calibrated to [Ca^2+^]_c_ from (Grynkiewicz, Poenie and Tsien, 1985):

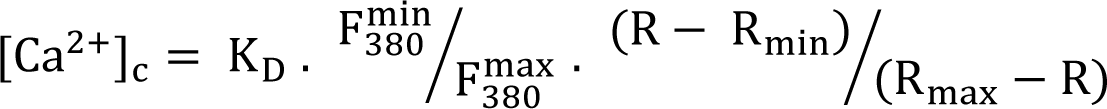

where, K_D_ = 225 nM (Forostyak *et al*., 2013).

### Western Blots

Proteins were isolated in RIPA buffer (Sigma, Cat# R0278) with protease inhibitor cocktail (Sigma, Cat# P8340) or, for WB of Orai1, in medium containing 150 mM NaCl, 50 mM Tris, 1% Triton-X-100, 0.1% SDS and protease inhibitor cocktail. After 30 min on ice with intermittent shaking, samples were collected by centrifugation (11,000 ×*g*, 20 min) and their protein content was determined (Thermo Pierce BCA Protein Assay kit, ThermoFisher, Cat# 23225). Proteins (∼30 µg/lane) were separated on 8% SDS-PAGE gels for IP_3_R or 10% SDS-PAGE gels for STIM1 and Orai1, and transferred to a Protran 0.45-μm nitrocellulose membrane (Merck, Cat# GE10600003) using a TransBlot semi-dry transfer system (BioRad, Cat# 1703940). Membranes were blocked by incubation (1 hr, 20°C) in TBST containing skimmed milk or bovine serum albumin (5%, Sigma, Cat# A9418). TBST (Tris-buffered saline with Tween) comprised: 137 mM NaCl, 20 mM Tris, 0.1% Tween-20, pH 7.5. Membranes were incubated with primary antibody in TBST (16 hr, 4°C), washed with TBST (3 ×10 min), incubated (1 hr, 20°C) in TBST containing HRP-conjugated secondary antibody (1:3000 anti-mouse, Cell Signaling Technology Cat# 7076S; or 1:5000 anti-rabbit, ThermoScientific Cat# 32260). After 3 washes, HRP was detected using Pierce ECL Western Blotting Substrate (ThermoFisher, Cat# 32106) and quantified using ImageQuant LAS 4000 (GE Healthcare) and Image J. The primary antibodies used were to: IP_3_R1 (1:1000, ThermoFisher, Cat# PA1-901, RRID: AB_2129984); β-actin (1:5000, BD Biosciences, Cat# 612656, RRID: AB_2289199); STIM1 (1:1000, Cell Signaling Technology, Cat# 5668S, RRID: AB_10828699); Orai1 (1:500, ProSci, Cat# PM-5205, RRID: AB_10941192); IP_3_R2 (1:1000, custom made by Pocono Rabbit Farm and Laboratory) (Mataragka and Taylor, 2018); and IP_3_R3 (1:500, BD Biosciences, Cat# 610313, RRID: AB_397705).

### Immunocytochemistry

After appropriate treatments, cells on a coverslip-bottomed plate were washed twice with cold PBS, fixed in PBS with paraformaldehyde (4%, 20°C, 20 min), washed (3 × 5 min) with PBS containing Triton-X100 (0.1%, PBST) and blocked by incubation (1 hr, 20°C) in PBST containing goat serum (5%). After incubation with primary antibody in PBST (16 hr, 4°C) and washing with PBST (3 × 5 min), cells were incubated (1 hr, 20°C) with secondary antibody in PBST containing goat serum, washed (3 × 5 min), stained (10 min, 20°C) with DAPI (1 µg/ml in PBS; Sigma, Cat# D9542) and washed (5 min, PBST). Cells were then covered with glycerol (60% v/v) and imaged using an Olympus FV300 confocal laser scanning microscope with 20× or 60× oil-immersion objectives. Fluorescence was analyzed using ImageJ. The primary antibodies used were to: PAX6 (1:500, Abcam, Cat# ab195045, RRID: AB_2750924); Nestin (1:500, Abcam, Cat# 92391, RRID: AB_10561437); Ki67 (1:250, Abcam, Cat# ab16667, RRID: AB_302459); SOX1 (1:1000, Abcam, Cat# ab87775, RRID: AB_2616563); Tuj1 (βIII Tubulin 1:1000, Promega, Cat# G712, RRID: AB_430874); NeuN (1:300, Abcam, Cat# ab177487, RRID: AB_2532109); Doublecortin (1:500, Abcam, Cat# 18723, RRID: AB_732011); MAP2 (1:200, Abcam, Cat# ab32454, RRID: AB_776174); STIM1 (1:1000, Cell Signaling Technology, Cat# 5668S, RRID: AB_10828699); and Orai1 (1:500, ProSci, Cat# PM-5205, RRID: AB_10941192).

### Proximity Ligation Assay

The Duolink *In Situ* Red Starter Mouse/Rabbit kit was from Sigma (#Cat DUO92101) and used according to the manufacturer’s protocol with primary antibodies to Orai1 (mouse 1:500) and STIM1 (rabbit 1:1000). Cells (∼30% confluent) were treated with thapsigargin (1 μM, 5 min) in Ca^2+^-free HBSS before fixation, permeabilization, and incubation with primary antibodies (16 hr, 4°C) and the PLA reactants. Red fluorescent PLA signals were imaged using an Olympus FV300 confocal laser scanning microscope, with excitation at 561 nm, and a 60× oil-immersion objective. Quantitative analysis of the intensity and surface area of PLA spots used the “Analyze particle” plugin of Fiji. Results are shown for 8-10 cells from two biological replicates of each genotype.

### Detection of STIM1 and IP_3_R1 puncta using TIRF microscopy

SHSY5Y cells were cultured on 15 mm glass coverslips coated with poly-D-lysine (100 μg/ml) in a 35 mm dish for 24 h. Cells were co-transfected with 500 ng of mCherry rIP_3_R1 and 200 ng mVenus STIM1 plasmids using TransIT-LT1 transfection reagent in Opti-MEM. Following 48 h of transfection and prior to imaging, cells were washed with imaging buffer (10 mM HEPES, 1.26 mM Ca^2+^, 137 mM NaCl, 4.7 mM KCl, 5.5 mM glucose, 1 mM Na_2_HPO_4_, 0.56 mM MgCl_2_, at pH 7.4). The coverslips were mounted in a chamber and imaged using an Olympus IX81 inverted total internal reflection fluorescence microscope (TIRFM) equipped with oil-immersion PLAPO OTIRFM 60× objective lens/1.45 numerical aperture and Hamamatsu ORCA-Fusion CMOS camera. Olympus CellSens Dimensions 2.3 (Build 189987) software was used for imaging. The angle of the excitation beam was adjusted to achieve TIRF with a penetration depth of ∼130 nm. Images were captured from a final field of 65 µm × 65 µm (300 × 300 pixels, one pixel=216 nm, binning 2×2). Cells positive for both mCherry rIP_3_R1 and mVenus STIM1 were identified using 561 nm and 488 nm lasers, respectively. The cells were incubated in zero calcium buffer (10 mM HEPES, 1 mM EGTA, 137 mM NaCl, 4.7 mM KCl, 5.5 mM glucose, 1 mM Na2HPO4, 0.56 mM MgCl2, at pH 7.4) for 2 minutes followed by addition of 30 µM CPA in zero calcium buffer. IP_3_R1 and STIM1 puncta prior to CPA addition and after CPA addition were captured at 1 min intervals. Raw images were filtered for background correction and same setting was used across all samples. Regions where fresh STIM1 puncta (0.2- 10 pixels) appeared post-CPA treatment at 10 mins were marked and subsequently IP_3_R1 puncta (0.2-10 pixels) were captured from the same region. Particle analysis and RGB profile plot were done using ImageJ.

### Statistical Analyses

All experiments were performed without blinding or prior power analyses. Independent biological replicates are reported as the number of experiments (n), with the number of cells contributing to each experiment indicated in legends. The limited availability of materials for PLA restricted the number of independent replicates (n) to 2 (each with 8-10 cells). Most plots show means ± s.e.m. (or s.d.). Box plots show 25^th^ and 75^th^ percentiles, median and mean (see legends). Where parametric analyses were justified by a Normality test, we used Student’s *t*-test with unequal variances for 2-way comparisons and ANOVA followed by pair-wise Tukey’s test for multiple comparisons. Non-parametric analyses used the Mann-Whitney U-test. Statistical significance is shown by \*\*\**P* < 0.001, \*\**P* < 0.01, \**P* < 0.05, or by letter codes wherein different letters indicate significantly different values (*P* < 0.001, details in legends). All analyses used Origin 8.5 software.

### Details of the plasmids and recombinant DNAs are given in Supplementary file 1

#### Source data files

Figure 1 source data

Figure 2 source data

Figure 3 source data

Figure 4 source data

Figure 5 source data

Figure 6 source data

Figure 7 source data

Figure 1 – figure supplement 1 source data

Figure 2 – figure supplement 1 source data

Figure 2 – figure supplement 2 source data

Figure 2 – figure supplement 3 source data

Figure 3 – figure supplement 1 source data

Figure 4 – figure supplement 1 source data

Figure 6 – figure supplement 1 source data

