## Supplemental file 1 for "Regulation of Store-Operated Ca^2+^ Entry by IP_3_ Receptors Independent of Their Ability to Release Ca^2+^"

| **Recombinant DNA** | | |
| --- | --- | --- |
| Lentiviral pZIP transfer vectors encoding non-silencing shRNA (NS, NT#3-TTGGATGGGAAGTTCACCCCG) | TransOMIC Technologies, Huntsville, AL | Custom made |
| Lentiviral pZIP transfer vectors encoding IP_3_R1-targeting shRNA (ULTRA3316782- TTTCTTGATCACTTCCACCAG) | TransOMIC Technologies | Custom made |
| pCMV- dR8.2 dpvr plasmid | Addgene, Watertown, MA | Cat #8455 |
| pCMV-VSV-G | Addgene | Cat #8454 |
| Human Orai1 siRNA | Dharmacon, Lafayette, CO | Cat# L-014998-00-0005 |
| Non-silencing (NS) siRNA (control for Orai1 siRNA) | Dharmacon | Cat# D-495 001810-10-05 |
| Human STIM1 siRNA | Santa Cruz Biotechnology Inc,  Dallas, TX | Cat# sc-76589 |
| Non-silencing (NS) siRNA (control for STIM1 siRNA) | Santa Cruz Biotechnology | Cat# sc-37007 |
| IP_3_R1 (rat type IP_3_R1 in pcDNA3.2/V5DEST vector) | Dellis et al., 2008 |  |
| Rat IP_3_R1^DA^ (D2550 replaced by A in IP_3_R1 in pcDNA3.2 vector) | Dellis et al., 2008 |  |
| Rat IP_3_R1^RQ^ (R568 replaced by Q of IP_3_R1 in pCDNA3.2/V5DEST vector) | Dellis et al., 2008 |  |
| Rat IP_3_R1^RQ/KQ^ (R568 and K569 replaced by Q of IP_3_R1 in pCDNA3.2/V5DEST vector) | Dellis et al., 2008 |  |
| Rat IP_3_R1^1-604^ (residues 1-604 of IP_3_R1 with N-terminal GST tag in pcDNA3.2/V5DEST vector) | Dellis et al., 2008 |  |
| Rat IP_3_R3 (rat IP_3_R3 in pcDNA3.2/V5DEST vector) | Saleem et al., 2013 |  |
| Human mCherry-STIM1 (N terminal mCherry tagged human STIM1 in pENTR1a vector) | Nunes-Hasler et al., 2017 |  |
| Human extended synaptotagmin 1 (E-Syt1) | Gift from Prof. S. Muallem, NIDCR, USA | Maléth et al., 2014 |
| Cas9n expressing plasmid (pSpCas9n(BB)-2A-Puro (PX462) V2.0) | Addgene | Plasmid #62987 |
| IP_3_R1 exon 3 targeted sgRNA forward (5’ CACCGCATTTGTTCTCTGTACGCGG 3’) | Eurofin | Custom made |
| IP_3_R1 exon 3 targeted sgRNA reverse (5’ AAACCCGCGTACAGAGAACAAATGC 3’) | Eurofin | Custom made |
| sgRNA validation primer for genomic PCR  FWD- 5’ TGTCTAGCTTCCTACATATTGGAGA 3’  REV – 5’ AACAAACCGTGCCACACAAG 3’ | Eurofin | Custom made |
| Reference gene GAPDH primers for genomic DNA PCR  FWD- 5’ TCACCAGGGCTGCTTTTAACTC 3’  REV- 5’ ATGACAAGCTTCCCGTTCTCAG 3’ | Sigma Aldrich | Custom made |
| sgRNA validation primer for Droplet Digital PCR  FWD- 5’ TCCCCTTCTGAACATTTCTTTTCT 3’  REV- 5’ TGCATGCTCTTACCCCAAGG 3’ | Eurofin | Custom made |
| Reference gene RPP30 primer for Droplet Digital PCR  FWD- 5’ AAGAAAGCCAAGTGTGAGGG 3’  REV- 5’ GGAAGAAGGGAGTGCTGACA 3’ | Eurofin | Custom made |
| mCherry-rat IP_3_R1 ^R568Q/K569Q^ (mCherry-rat IP_3_R1^RQ/KQ^)  FWD: 5’GCAAGACTACcagcagAACCAGGAGTAC-3’  REV:  5’-TGTGAGTGTCTCAGGACC-3’ | IDT | Custom made |
| **Software and algorithms** | | |
| AndoriQ imaging software version 2.4.2 | Andor | RRID: SCR_014461 |
| ImageQuant LAS 4000 software version 1.2 | GE Healthcare | RRID: SCR_014246 |
| ImageJ version 1.52d | ImageJ | RRID: SCR_003070 |
| Fiji version 1.52p | ImageJ | RRID: SCR_002285 |
| Olympus FluoView software FV31S-SW version 2.3.1 | Olympus LS |  |
| Origin 8.5 software | OriginLab Corporation | RRID: SCR_014212 |
| QuantaSoft version 1.7.4.0917 | Bio-Rad |  |
| CRISPOR | Tefor infrastructure | http://crispor.tefor.net/ |
| Benchling | Benchling | https://benchling.com/ |
| Primer3web version 4.1.0 | ELIXIR | https://primer3.ut.ee/ |
| BioRender | BioRender | RRID: SCR_018361 |
